# Global and context-specific transcriptional consequences of oncogenic Fbw7 mutations

**DOI:** 10.1101/2021.10.19.464955

**Authors:** H. Nayanga Thirimanne, Feinan Wu, Derek H Janssens, Jherek Swanger, Heather M Feldman, Robert A Amezquita, Raphael Gottardo, Patrick J Paddison, Steven Henikoff, Bruce E Clurman

## Abstract

Fbw7 is a ubiquitin ligase substrate receptor that targets proteins for proteasomal degradation. Most known Fbw7 substrates are transcription factors (TFs) and many are also oncoproteins (e.g., c-Myc, c-Jun, Notch). Fbw7 is an important tumor suppressor and *FBXW7* mutations drive tumorigenesis through activation of oncogenic Fbw7 substrates. Defining the mechanisms of Fbw7-associated tumorigenesis is critical for developing targeted therapies. We thus determined the transcriptional consequences of oncogenic Fbw7 mutations by studying isogenic colorectal cancer cell lines with engineered *FBXW7* null and heterozygous missense mutations. We used an integrated approach employing RNA-Seq and high-resolution mapping (CUT&RUN) of histone modifications and TF occupancy (c-Jun and c-Myc) to examine the combinatorial effects of mis-regulated Fbw7 substrates. Fbw7 mutations caused widespread transcriptional changes associated with active chromatin and altered TF occupancy at distal regulatory regions. Some regulatory changes were common to both *FBXW7*-mutant cell lines whereas others were *FBXW7* mutation-specific. By comparing c-Jun and c-Myc binding sites, we also identified co-regulated elements, suggesting that Fbw7 substrates may have synergistic effects. One co-regulated gene was CIITA, a master regulator of MHC Class II gene expression, and Fbw7 loss increased CIITA and MHC Class II gene expression in colorectal cancer cells. Fbw7 mutations were also correlated with increased CIITA expression in TCGA colorectal tumors and cell lines, which may have immunologic implications for progression and treatment of Fbw7-associated cancers. This integrative analysis provides a framework for understanding normal and neoplastic context-specific Fbw7 functions.

## Introduction

SCFs (Skp1-Cul1-F-box protein) are multi-subunit ubiquitin ligases that target proteins for degradation through the conjugation of polyubiquitin chains that signal their destruction by the proteasome (Deshaies & Joazeiro, 2009; Lee & Diehl, 2014). F-box proteins are SCF substrate receptors and often target proteins for ubiquitylation in response to substrate modifications (Skaar et al., 2013; Yumimoto & Nakayama, 2020). The Fbw7 F-box protein is encoded by the *FBXW7* gene and binds to substrates after they become phosphorylated within motifs, termed CDC4-phosphodegrons (CPDs), that mediate high-affinity interactions with the Fbw7 β-propeller (R. J. Davis et al., 2014; Hao et al., 2007; Nash et al., 2001; Orlicky et al., 2003; Yumimoto & Nakayama, 2020). Fbw7 also contains a dimerization domain and an F-box that binds to the SCF complex, and Fbw7 brings phosphorylated substrates into proximity with the remainder of the SCF complex.

Approximately 30 Fbw7 substrates are known. Most are broadly acting transcription factors (TFs) that control processes such as proliferation, differentiation, and metabolism, and include c-Myc, Notch, c-Jun, PGC-1α, SREBP1/2, and many others (Cremona et al., 2016; R. J. Davis et al., 2014; Welcker & Clurman, 2008; Yumimoto & Nakayama, 2020). Fbw7 also targets proteins that are not TFs, such as cyclin E and MCL-1. Fbw7’s cellular functions reflect the combined regulation of its many substrates. Because different cell types express different subsets of substrates that are targeted for degradation by Fbw7 only after they acquire specific phosphorylations, the contribution of individual substrates to Fbw7’s overall function is highly context dependent. In most known cases, glycogen synthase kinase 3β (GSK-3β) is one of the CPD kinases, suggesting that Fbw7 coordinately couples the activity of its substrates to mitogenic signaling pathways. In addition, some TFs are phosphorylated when they are associated with their target genes (Fryer et al., 2004; Punga et al., 2006), highlighting another constraint to their susceptibility to Fbw7-mediated degradation. This complexity has made it difficult to fully comprehend Fbw7 function, and this is compounded by the fact that many substrates are master TFs that regulate complex gene networks themselves.

Some Fbw7 substrates are critical oncoproteins that drive tumorigenesis, such as c-Myc, c-Jun, and Notch1. Fbw7 loss in tumors deregulates these oncoproteins and *FBXW7* is a commonly mutated tumor suppressor gene (R. J. Davis et al., 2014; Shimizu et al., 2018; Tan et al., 2008; Yeh et al., 2018; Yumimoto & Nakayama, 2020). The most frequent *FBXW7* mutations are heterozygous missense mutations, termed Fbw7^R/+^, that target one of three arginine residues that form Fbw7’s CPD-binding pocket. Fbw7^R/+^ weaken substrate binding and are thought to act as dominant negatives by forming heterodimers with WT-Fbw7, which normally functions as a dimer (Hao et al., 2007; Welcker et al., 2013; Welcker & Clurman, 2007). While Fbw7^R/+^ mutations are common, Fbw7^+/-^ mutations are not, strongly suggesting that Fbw7^R/+^ mutations are not simply loss-of-function alleles and that Fbw7^R/+^ proteins have unique oncogenic activity (R. J. Davis et al., 2014). However, the mechanisms that drive Fbw7^R/+^ selection in cancers are still poorly understood. One model, termed “just enough”, posits that Fbw7^R/+^ only partially impair Fbw7 and inactivate its tumor suppressor functions but preserve other required or beneficial Fbw7 activities (H. Davis & Tomlinson, 2012). Unlike Fbw7^+/-^ mutations, canonical bi-allelic loss of function Fbw7^-/-^ mutations (e.g., nonsense, truncations, frame shifts, deletions) do occur. Different cancers have different mutational spectra; T-cell acute lymphocytic leukemias (T-ALLs) have almost exclusively Fbw7^R/+^ whereas colorectal cancers have both Fbw7^R/+^ and Fbw7^-/-^ mutations. The mechanisms through which these different mutations promote tumorigenesis remain poorly defined, yet this is critical to understanding tumor suppression by Fbw7 and developing targeted therapeutic strategies.

To address these questions, we determined the global transcriptional consequences of oncogenic Fbw7 mutations by using RNA-Seq and high-resolution mapping of histone modifications and oncogenic TF (c-Jun and c-Myc, here onwards Jun and Myc) occupancy in an isogenic panel of colorectal cancer cells with engineered Fbw7^-/-^ and Fbw7^R/+^ mutations. Both mutations caused widespread but highly context-specific transcriptional changes associated with active chromatin and altered TF occupancy. Overall, the transcriptional effects of Fbw7^-/-^ were greater than Fbw7^R/+^. We found evidence supporting the “just enough” model of Fbw7^R/+^ (sites shared by both mutations, but less impacted by Fbw7^R/+^), as well as outcomes specific to either the Fbw7^-/-^ or Fbw7^R/+^ mutation.

While both mutations only impacted a subset of mapped loci, we found sites at which both Jun and Myc were co-regulated by Fbw7. One co-regulated gene was *CIITA* (Class II Major Histocompatibility Complex Transactivator), the master regulator of MHC class II gene expression (Masternak et al., 2000; Reith et al., 2005). Fbw7 loss caused increased MHC class II gene expression associated with increased Jun and Myc occupancy upstream of *CIITA*. Analyses of TCGA colorectal cancer and cell lines further correlated Fbw7 mutations with MHC Class II gene expression, which may have important prognostic and therapeutic implications for Fbw7-associated colorectal cancers. Because Fbw7 normally regulates neural stem cells (NSCs) (Hoeck et al., 2010) and Fbw7 loss occurs in glioblastoma (Hagedorn et al., 2007), we similarly studied NSCs in which Fbw7 was acutely deleted, which revealed transcriptional consequences of Fbw7 loss that closely mirrored the results obtained in the isogenic cell panel, including increased CIITA expression. Overall, these data establish a framework for understanding the mechanisms of Fbw7 tumor suppression.

## Results

### *FBXW7* null and missense mutations lead to distinct gene expression profiles

Hct116 cells were previously engineered to mutate the endogenous wild-type (WT) *FBXW7* locus to either a heterozygous Fbw7^R505C/+^ (Fbw7^R/+^) or a homozygous null (Fbw7^-/-^) mutation (Figure 1A) (R. J. Davis et al., 2018; Grim et al., 2008). We performed RNA sequencing to identify the global transcriptome changes arising in response to these Fbw7 mutations. Principal component analysis (PCA) revealed that the Fbw7^R/+^ and Fbw7^-/-^ cells clustered apart from one another, indicating that the two *FBXW7* mutations have distinct effects on the transcriptome relative to WT cells (Figure 1 – figure supplement 1). Compared with WT cells, 11.3% and 5.4% of protein-coding genes were differentially expressed in Fbw7^-/-^ and Fbw7^R/+^ cells, respectively. Some genes were differentially expressed in both Fbw7^-/-^ and Fbw7^R/+^, whereas others were uniquely deregulated by either Fbw7^-/-^ or Fbw7^R/+^ (Figure 1B, Figure 1 – source data 1). Hierarchical clustering of differentially expressed protein-coding genes identified genes that were: 1) upregulated (cluster 1) or downregulated (cluster 2) in just Fbw7^-/-^ cells, 2) genes upregulated (cluster 6) and downregulated (cluster 5) in just Fbw7^R/+^ cells, and 3) genes that show similar expression changes in response to both *FBXW7* mutations (clusters 3 and 4) (Figure 1C). Gene set enrichment analysis revealed numerous pathways enriched in the differentially expressed genes common to both types of Fbw7 mutations or uniquely to Fbw7^-/-^ cells (Figure 1D) (E. Y. Chen et al., 2013; Kuleshov et al., 2016). The most highly enriched Gene Ontology (GO) term in Fbw7^-/-^ cells was Major Histocompatibility Complex Class II (MHC Class II) components. Some enriched terms might reflect the functions of known Fbw7 substrates, such as the regulation of cholesterol biosynthesis by the SREBP-1/2 proteins (Sundqvist et al., 2005). The two distinct *FBXW7* mutations thus caused both overlapping and unique changes in global transcription.

**Figure 1.**
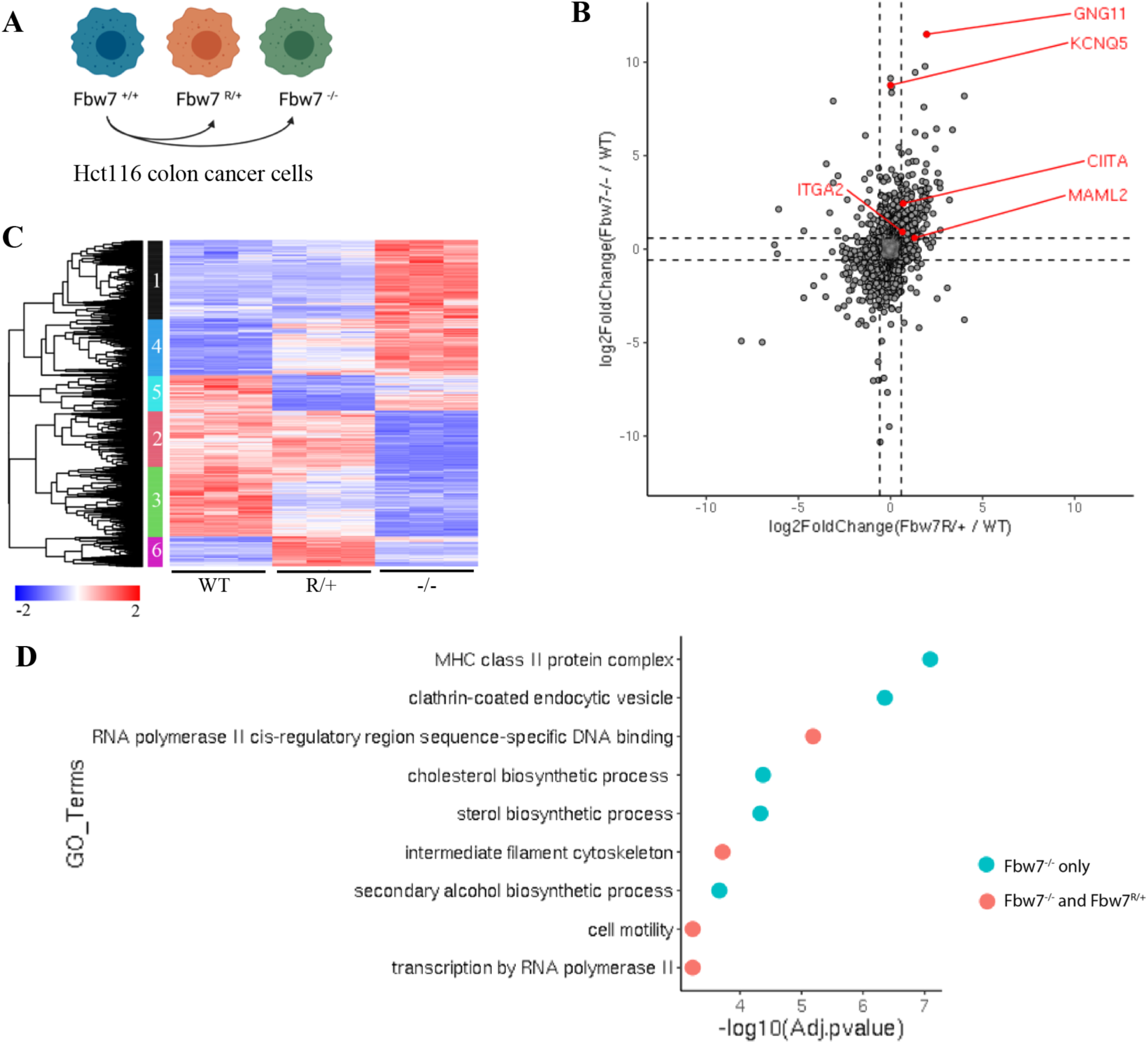
RNA-Seq reveals differential gene expression in Hct116 Fbw7^-/-^ and Fbw7^R/+^ cells. (A) Genetically engineered isogenic cell lines used in the study: Hct116 wild-type (WT), Fbw7^-/-^ and Fbw7^R/+^. (B) Differentially expressed protein-coding genes (represented by each dot) in Fbw7^-/-^ or Fbw7^R/+^ (FDR<0.05). Dashed lines mark log_2_FC=0.6. Three replicates per cell type were included. (C) Hierarchical clustering of differentially expressed protein-coding genes. The heatmap shows the intensity of expression of each gene (y axis) for three replicates per cell type (x axis). (D) Gene ontology terms that are enriched in gene clusters deregulated by both Fbw7^-/-^ and Fbw7^R/+^ (red), and only by Fbw7^-/-^ (blue). Detailed output of differential expression analysis, hierarchical clustering and GO Term analysis are provided as Figure 1-source data file 1, 2 and 3 respectively. See Figure 1 – figure supplement 1 for the PCA of Hct116 RNA-Seq.

### Chromatin regulation in *FBXW7* mutant cells

Because Fbw7 regulates many TFs, we first looked at global changes in chromatin to determine if specific TFs targeted by Fbw7 might drive the transcriptional changes in these cells. Histone H3 lysine-27 acetylation (H3K27ac) and Histone H3 lysine-27 trimethylation (H3K27me3) provide a simple readout of transcriptionally active versus repressive chromatin, respectively (Karlić et al., 2010). We used Cleavage Under Target and Release Using Nuclease (CUT&RUN) (Janssens et al., 2018; Skene et al., 2018; Skene & Henikoff, 2017) to obtain high resolution maps of H3K27ac and H3K27me3 in each of the Hct116 cell lines (Figure 2 – figure supplement 1). As expected, the H3K27ac signal within the 2 kb region flanking the transcriptional start sites (TSSs) of genes was positively correlated with their expression (r = 0.44, p value < 2.2e-16), whereas the amount of H3K27me3 was negatively correlated (r = -0.22, p value < 2.2e-16) (Figure 2A). For example, the *GNG11* gene, whose expression is upregulated in Fbw7^-/-^ cells, contains increased H3K27ac and decreased H3K27me3, compared with WT (Figure 1B, Figure 2B)

**Figure 2.**
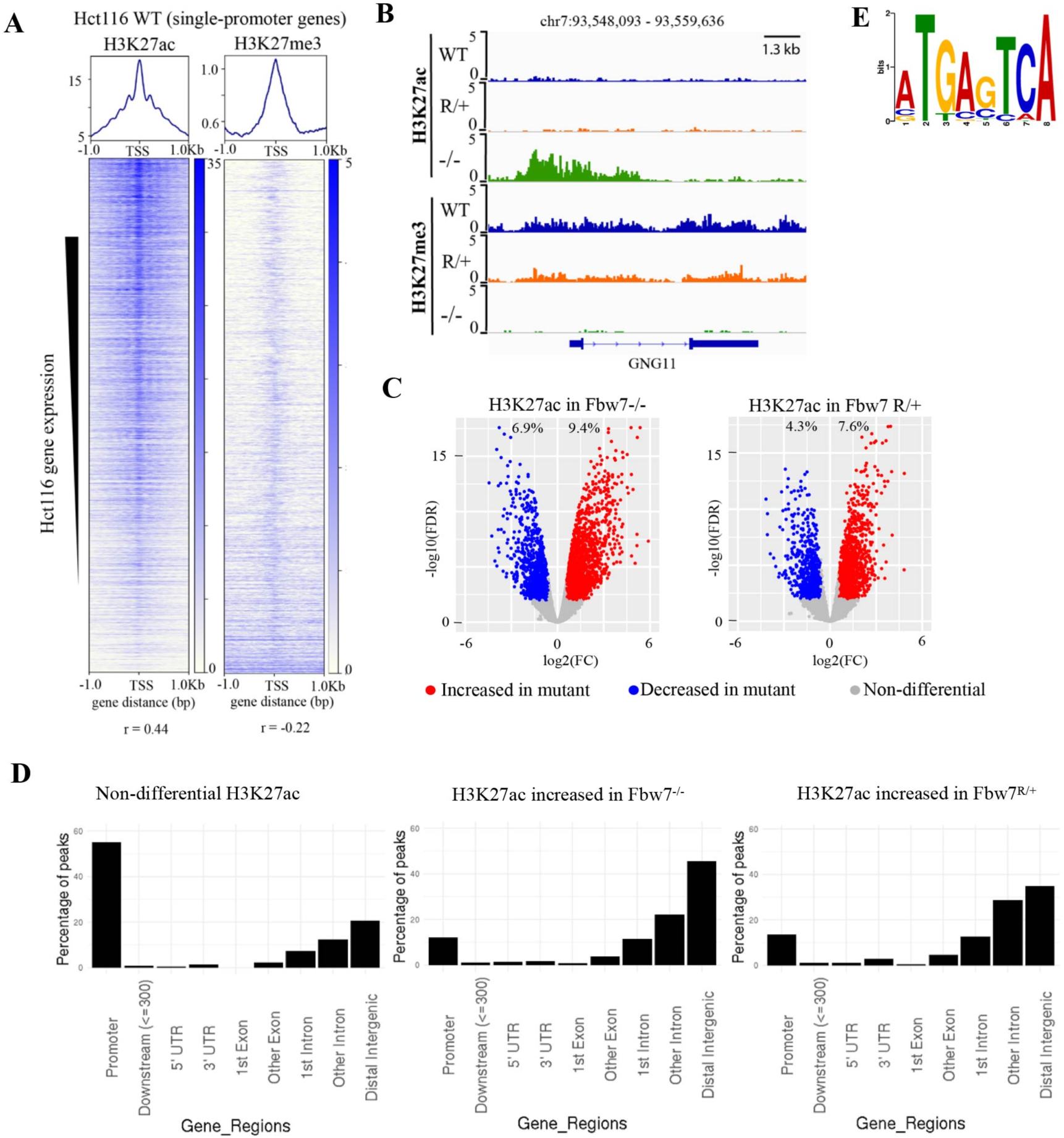
Differential H3K27ac signal in Hct116 *FBXW7* mutant cells reveals genomic sites targeted by Fbw7. (A) Heatmaps showing the correlation between CUT&RUN profiles of H3K27ac and H3K27me3, and RNA-Seq in Hct116 WT cells. (B) Genome browser view of H3K27ac and H3K27me3 signal from Hct116 WT, Fbw7^R/+^ and Fbw7^-/-^ cells at a representative gene (C) Peaks with increased (red) or decreased (blue) H3K27ac signal in Hct116 Fbw7^-/-^ and Fbw7^R/+^ cells compared to WT cells. Differential sites indicated as a percent of total H3K27ac peaks in Hct116 WT cells. See Figure 2 – figure supplement 1 for the correlation matrix, Figure 2 – source data 1 for a list of H3K27ac differential sites and Figure 2 – source data 2 for the calculation of percentages. (D) Percentage of H3K27ac peaks located within different gene regions. See Figure 2 – figure supplement 2. (E) Sequence logo for AP-1 motif enriched in H3K27ac peaks increased in Fbw7^-/-^ cells (E value = 1.6e-3). See Figure 2 – figure supplement 3 for the complete MEME output and details on the FIMO analysis.

Genome-wide analysis identified sites with increased H3K27ac in *FBXW7* mutant cells (Fbw7^-/-^: 9.4%, Fbw7^R/+^: 7.6%) compared with control cells, as well as sites where H3K27ac was decreased (Fbw7^-/-^: 6.9%, Fbw7^R/+^: 4.3%) (Figure 2C, Figure 2 - source data 2). Most non-differential H3K27ac sites (those unaffected by Fbw7 status) were promoter-proximal, while loci with differential H3K27ac in either Fbw7^R/+^ or Fbw7^-/-^ cells fell mostly within introns or intergenic regions (p value < 0.0001, Fisher test) (Figure 2D, Figure 2 – figure supplement 2). To determine whether these differential loci result from the altered binding of known Fbw7 substrates, we performed motif discovery analysis on the central 100 bp sequence of each peak. Strikingly, the AP-1 motif, which is bound by the Jun family, was found in 32% (p value ≤ 1.8e-5) of the H3K27ac sites upregulated in Fbw7^-/-^ cells (Figure 2E, Figure 2 – figure supplement 3A). The AP-1 motif was also enriched in differential H3K27ac sites that were decreased in Fbw7^-/-^ cells, as well as in differential H3K27c sites in Fbw7^R/+^ cells. In contrast, the AP-1 site was not enriched in H3K27ac sites that were unaffected by either *FBXW7* mutation (Figure 2 – figure supplement 3B). AP-1 motif enrichment in these differential sites suggests that Fbw7-dependent Jun regulation may account, in part, for these changes.

### Fbw7 preferentially regulates Jun and Myc occupancy at distal regulatory regions

Fbw7 targets some TF-substrates while they are bound to DNA, suggesting that substrates may recruit Fbw7 to chromatin (Fryer et al., 2004; Punga et al., 2006). We thus examined how mutation of the Fbw7 substrate binding domain impacts its chromatin association in Hct116 cells with endogenous heterozygous (Fbw7^R/+^) or homozygous (Fbw7^R/R^) mutations. Fbw7 was found in both the chromatin and soluble fractions of WT-Hct116 lysates, but exclusively in the soluble fraction in Fbw7^R/R^ cells (Figure 3A). The only known consequence of Fbw7^R^ mutations is to prevent substrate binding, and thus the loss of chromatin-associated Fbw7 in Fbw7^R/R^ cells suggests that substrate binding recruits Fbw7 to chromatin. Proteasome inhibition prevents substrate degradation and stabilizes Fbw7-substrate complexes. We found that treatment of cells with a proteasome inhibitor, bortezomib, further shifted Fbw7 to chromatin (Figure 3 – figure supplement 1), supporting the hypothesis that substrate binding underlies Fbw7 chromatin association.

**Figure 3.**
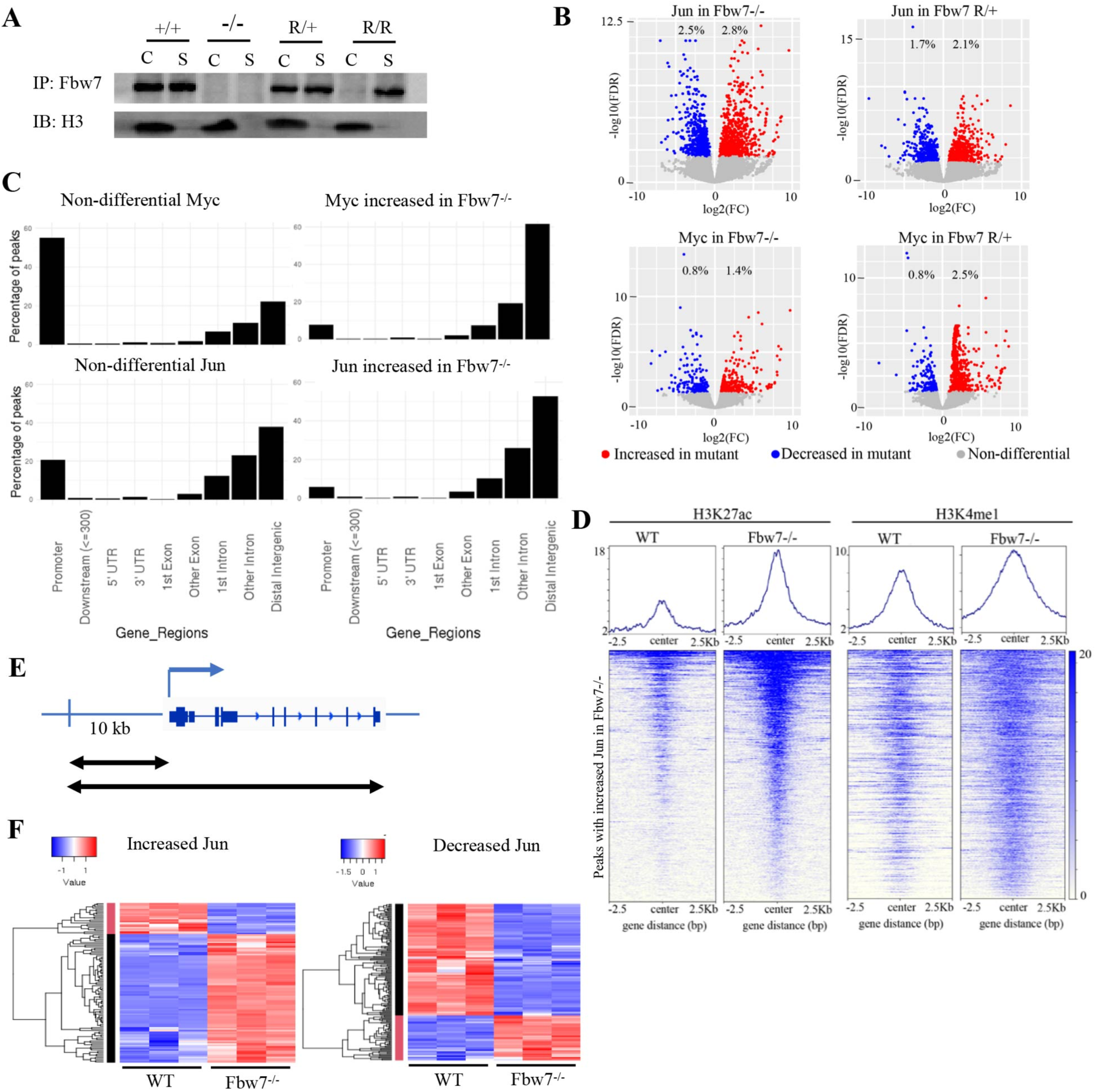
Fbw7 regulates the occupancy of Jun and Myc on DNA, preferentially at distal regulatory regions. (A) Fbw7 abundance in chromatin (C) and soluble (S) fractions from Hct116 WT, Fbw7^R/+^ and Fbw7^R/R^ cells. Histone H3 was detected in chromatin fractions. (B) Increased (red) and decreased (blue) Jun and Myc sites in Hct116 Fbw7^-/-^ and Fbw7^R/+^ cells compared to WT. (C) Non-differential and differential Jun and Myc peaks located within gene features. (D) H3K27ac and H3K4me1 CUT&RUN signal from Hct116 WT and Fbw7^-/-^ cells mapped on genomic sites that have increased Jun occupancy in Fbw7^-/-^ cells. (E) Schema depicting the filtering criteria applied to the annotated differential sites to select gene proximal sites. (F) Transcription of genes that have increased or decreased Jun bound at a gene proximal site. (Each row is a gene and three replicates each from Hct116 WT and Fbw7^-/-^ cells are shown.) See Figure 3 – figure supplement 1-4 and Figure 3 – source data 1-2.

Myc and Jun are TF substrates with important roles in Fbw7-associated cancers (R. J. Davis et al., 2014). Myc deregulation in cancers results from either Fbw7 or Myc-CPD mutations (R. J. Davis et al., 2014; Welcker et al., 2004; Yada et al., 2004; Yumimoto & Nakayama, 2020). Myc stabilization is an important driver of Fbw7-associated tumorigenesis (R. J. Davis et al., 2014; O. M. Khan et al., 2018; King et al., 2013; Reavie et al., 2013; Yumimoto & Nakayama, 2020). CPD phosphorylation and Myc ubiquitylation also modulate Myc transcriptional activity (Endres et al., 2021; Gupta et al., 1993; Hemann et al., 2005; Jaenicke et al., 2016; Thomas & Tansey, 2011). Fbw7 loss stabilizes Myc in Hct116 cells but Myc steady state abundance is less impacted, due to Myc autoregulation (Grim et al., 2008). Fbw7 also targets Jun for degradation after multisite phosphorylation (Csizmok et al., 2018; Nateri et al., 2004; Wei et al., 2005) and several factors regulate Jun degradation by Fbw7 in Hct116 cells, including Rack1 (Zhang et al., 2012), BLM (Priyadarshini et al., 2018) and Usp28 (Diefenbacher et al., 2014).

We profiled genome-wide Jun and Myc occupancy (Figure 3 – figure supplement 2A) to determine the extent to which they are deregulated by Fbw7 mutations. As expected, Jun-binding and Myc-binding site motifs were highly enriched in the respective datasets (Figure 3 – figure supplement 2B). Differential binding analyses of the Jun and Myc peaks demonstrated that 5.3% and 3.8% of the Jun sites and 2.2% and 3.3% of the Myc sites exhibited differential occupancy in the Fbw7^-/-^ and Fbw7^R/+^ cells, respectively (Figure 3B, Figure 2 – source data 2). Thus, Fbw7 mutations lead to changes in Myc and Jun occupancy at specific loci, rather than a global increase in their chromatin occupancy.

Like H3K27ac, most non-differential Myc binding sites were promoter-proximal, whereas almost all differential sites with increased Myc occupancy in *FBXW7* mutant cells fell within introns and intergenic regions (p value < 0.001, Fisher test) (Figure 3C, Figure 3 – figure supplement 3). Compared with Myc, a smaller proportion of the total Jun sites in WT-Hct116 cells were promoter-proximal, but again the sites with differential occupancy in Fbw7 mutant cells were heavily biased to intron and intragenic regions (p value < 0.0001, Fisher test) (Figure 3C). The differential sites in introns and intergenic loci were also enriched for H3K27ac and H3K4me1, which is consistent with the idea that these sites function as distal regulatory elements, such as enhancers (Figure 3D).

To study the functional significance of Fbw7-dependent changes in Jun and Myc binding, we examined the expression of genes that could be linked to differential Jun or Myc sites (within the gene body or 10 kb upstream of TSS) (Figure 3E). Approximately 40% of genes with increased promoter-proximal Jun occupancy and 46% of genes with decreased promoter-proximal Jun occupancy in Fbw7^-/-^ cells exhibited corresponding increases or decreases in RNA expression (Figure 3F). Other genes linked to differential sites were either not captured by RNA-Seq or had expression changes that were statistically non-significant. Similar associations were seen with Myc differential sites, although fewer could be linked with transcripts than for Jun (Figure 3 – figure supplement 4). Overall, the differential sites that could be linked with associated genes showed strong concordance between the change in Jun or Myc occupancy and RNA expression.

### Fbw7^-/-^ and Fbw7^R/+^ mutation-specific consequences on Jun and Myc occupancy

We next examined how Jun occupancy is differentially affected by Fbw7^R/+^ and Fbw7^-/-^ mutations. Many of the differential Jun sites were common to both mutant cell lines: 48% of differential Jun sites in Fbw7^R/+^ (252/530; p value <0.0001, Fisher test) and 35% of differential Jun sites in Fbw7^-/-^ (252/715) (Figure 4A). Representative Jun peaks that are increased in Fbw7^-/-^ and/or Fbw7^R/+^ are shown in Figure 4B: a) Jun occupancy at *KCNQ5* intronic sites was increased only in Fbw7^-/-^ cells; b) in *ITGA2*, Jun occupancy was increased in both Fbw7^-/-^ and Fbw7^R/+^, but to an intermediate level in Fbw7^R/+^; and c) in *MAML2*, Jun occupancy was increased in Fbw7^R/+^ more highly than in Fbw7^-/-^.

**Figure 4.**
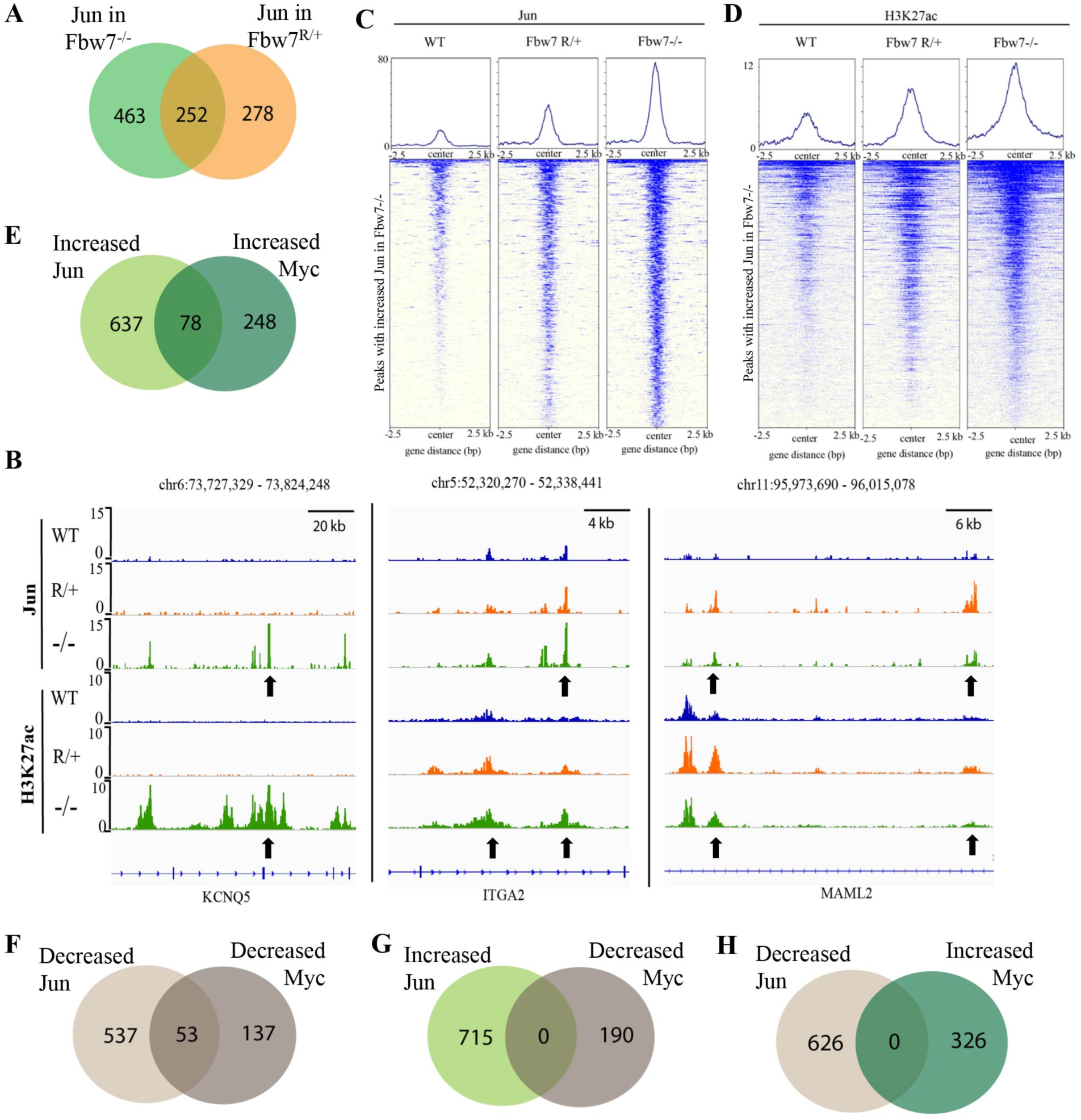
Fbw7 exhibits mutation-type specific regulation and coordinate regulation of multiple TFs. (A) The overlap between peaks with increased Jun in Fbw7^-/-^ and Fbw7^R/+^ cells. (B) Genome browser view of Jun and H3K27ac occupancy in Hct116 WT, Fbw7^-/-^ and Fbw7^R/+^ cells at representative loci. Black arrows point to peaks with increased signal uniquely in Fbw7^-/-^ (*KCNQ5*), in both Fbw7^-/-^ and Fbw7^R/+^ (intermediate level in Fbw7^R/+^) (*ITGA2*) and increased in Fbw7^R/+^ than in Fbw7^-/-^ (*MAML2*). (C, D) Heatmap of Jun and H3K27ac signal from each cell type mapped on sites with increased Jun in Fbw7^-/-^ cells. (E-H) E-The overlap between peaks with increased Jun and Myc, F- decreased Jun and Myc, G- increased Jun and decreased Myc, H-decreased Jun and increased Myc in Fbw7^-/-^ cells. See Figure 4 – figure supplement 1.

We found many sites like *ITGA2*, that exhibited an intermediate impact of Fbw7^R/+^ on Jun occupancy, compared with Fbw7^-/-^, as depicted by the heatmap of Jun signal from WT, Fbw7^R/+^, and Fbw7^-/-^ cells mapped on all the sites with increased Jun occupancy in Fbw7^-/-^ cells (Figure 4C). H3K27ac followed the same trend exhibited by Jun (Figure 4B, 4D). RNA-Seq data showed that genes in cluster 3 and 4 were deregulated in both Fbw7^-/-^ and Fbw7^R/+^, but to an intermediate level in Fbw7^R/+^ (Figure 1B, 1C). Because the Fbw7^-/-^ and Fbw7^R/+^ cells were derived independently, these shared sites and the gradients in Jun occupancy, H3K27ac, and RNA expression (Fbw7^-/-^ > Fbw7^R/+^) all support the conclusion that these differences are directly attributable to Fbw7 status.

We also identified Jun differential sites that were uniquely impacted by each mutation type (Figure 4A), which includes a subset of sites that were more strongly impacted by Fbw7^R/+^. RNA-Seq data also showed that genes in cluster 5 and 6 were deregulated most strongly in Fbw7^R/+^. In summary, we identified differential Jun sites that are uniquely affected by each Fbw7 mutation type and others that were shared between the two mutant cell lines.

### Fbw7 coordinately regulates Jun and Myc at co-occupied loci

Myc and Jun are oncogenic TFs with activities in shared pathways, and we next examined how they might be coregulated at shared sites. Approximately 20% of the Myc and Jun binding sites overlapped in Hct116 WT cells (Figure 4 – figure supplement 1A, p < 0.0001 Fisher Test) and Jun and Myc exhibited striking coordinate regulation by Fbw7 at these co-occupied differential loci. We identified 78 sites in which both Jun and Myc occupancy were increased in Fbw7^-/-^ cells and 53 sites where both Jun and Myc were decreased in Fbw7^-/-^ cells (Figure 4E, 4F). In contrast, no sites with discordant changes in Jun and Myc occupancy (e.g., increased Jun but decreased Myc) were found (Figure 4G, 4H). We found similar concordance in co-regulated Jun and Myc sites in Fbw7^R/+^ cells (Figure 4 – figure supplement 1B,1C).

### Jun and Myc co-regulation by Fbw7 controls MHC Class II gene expression

“MHC Class II” was the most enriched GO term in genes that exhibited altered expression in Fbw7^-/-^ cells (Figure 1C). Unlike the MHC class I genes, which are expressed in all cells, MHC Class II genes are normally expressed only in specific immune cells, and their expression is controlled by the Class II Major Histocompatibility Transactivator (CIITA) gene (Masternak et al., 2000; Ting & Trowsdale, 2002). We found that Myc and Jun occupancy were increased within *CIITA* upstream regulatory regions in Fbw7^-/-^ cells (Figure 5A).

**Figure 5.**
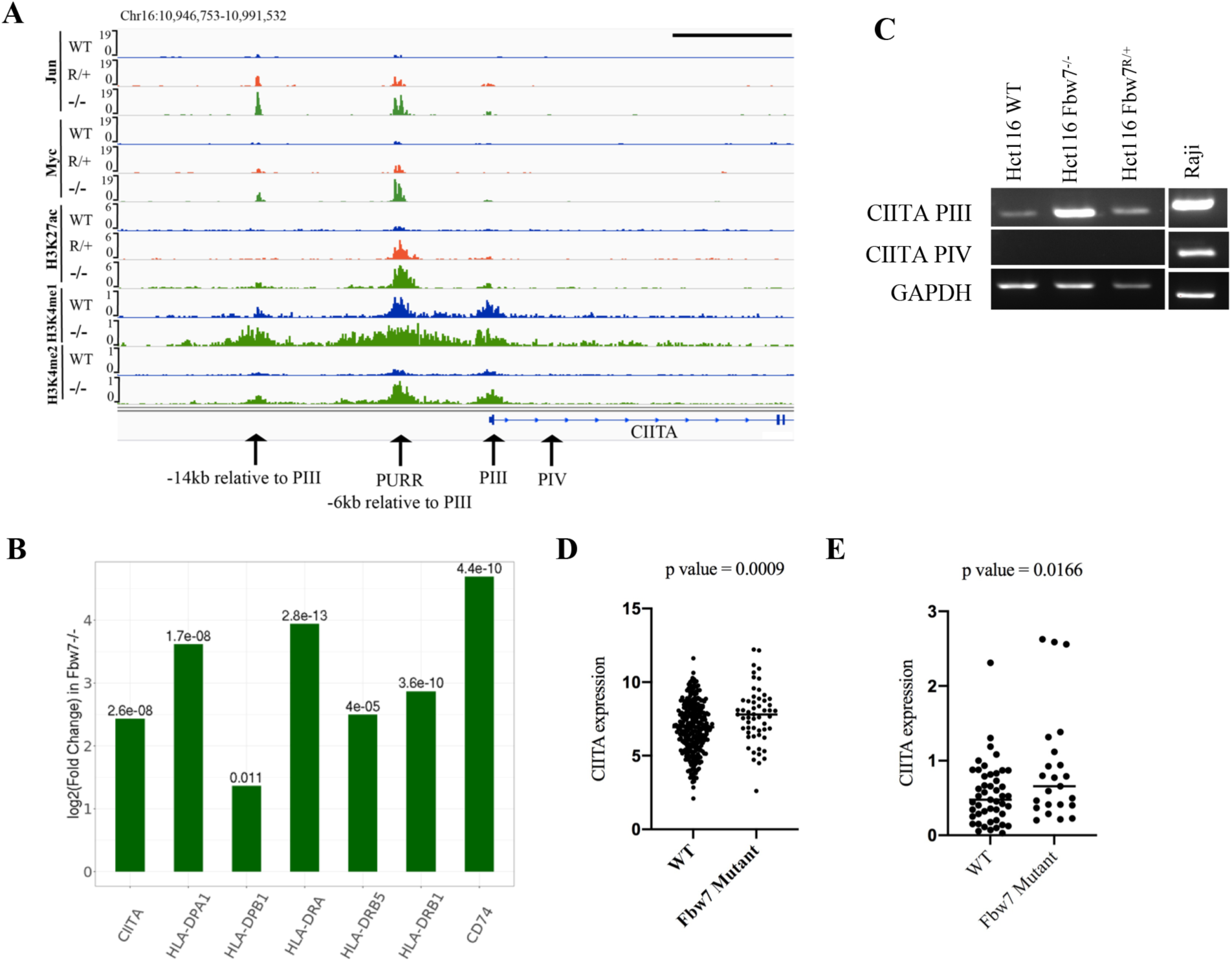
Fbw7 regulates the expression of MHC Class II genes. (A) Genome browser view of TFs and histone modification marks enriched at the promoter and regulatory sites upstream of *CIITA* gene. Arrows point to (from right to left): PIV (promoter of isoform IV); PIII (promoter of isoform III); PURR (PIII Upstream Regulatory Regions) - a known regulatory site 6 kb upstream of PIII; and a known regulatory site 14 kb upstream to PIII. Black scale bar = 10 kb (B) Expression fold change of CIITA and MHC Class II genes in Hct116 Fbw7^-/-^ with respect to WT cells. FDR values are indicated on top of each bar. n = 3 (C) CIITA isoform III and IV amplified using isoform specific primers in Hct116 and Raji cells. (D) CIITA expression in primary cancer samples from TCGA COADREAD datasets that have WT Fbw7 (n=297) and mutated Fbw7 (n=55). (E) CIITA expression in colon and rectal cancer cell lines with WT Fbw7 (n=47) and mutated Fbw7 (n=23). Data collected from DepMap portal. See Figure 5 – figure supplement 1, Figure 5 – source data 1-3

The *CIITA* gene contains four promoters (hereafter referred to as PI – PIV) that specify four transcripts with distinct first exons (Muhlethaler-Mottet et al., 1997). While CIITA isoform III is constitutively expressed in antigen presenting cells (APCs), isoform IV is inducible by cytokines in non-hematopoietic cells (van der Stoep et al., 2007). The PIII Upstream Regulatory Region (PURR) is located 6kb upstream of PIII and consists of regulatory sites for both constitutive and IFNγ-induced CIITA expression (Deffrennes et al., 2001; van der Stoep et al., 2007), as well as an AP-1 site (Martins et al., 2007). We found both Myc and Jun bound to regulatory elements (PURR and an element 14kb upstream of CIITA-PIII) and that their occupancy was increased in Fbw7^-/-^ cells (Figure 5A). Jun and Myc occupancy were also increased at these sites in Fbw7^R/+^ cells, but to a lesser extent. H3K27ac and H3K4me1 were increased at these sites in Fbw7^-/-^ cells, which is indicative of active transcription, and RNA-Seq revealed increased CIITA mRNA expression in Fbw7^-/-^ cells (Figure 5B). Isoform-specific primers demonstrated that the pIII isoform is elevated in Fbw7^-/-^ cells, but that the pIV isoform is not expressed (Figure 5C). Raji cells are shown as a control cell that expresses both CIITA isoforms. Importantly, the amount of upregulated CIITA expression in Fbw7^-/-^ cells is functionally significant and caused increased expression of MHC Class II genes that are targets of CIITA (HLA-DPA, HLA-DPB, HLA-DRB and HLA-DRA) (Figure 5B). In contrast, MHC Class I genes were not differentially expressed in *FBXW7* mutant cells (Figure 5 – figure supplement 1).

We also analyzed CIITA expression in primary colorectal cancers in TCGA datasets, which revealed increased CIITA expression in *FBXW7* mutant cancers compared with *FBXW7* WT tumors (Figure 5D). Because these primary tumors contain immune infiltrates, the increased CIITA expression could reflect CIITA expression in either tumor cells or immune cells. We thus analyzed colorectal cancer cell lines in the Cancer Cell Line Encyclopedia, which also exhibited elevated CIITA expression in *FBXW7* mutant cell lines (Figure 5E) (Ghandi et al., 2019). These data support the idea that Fbw7 regulates CIITA expression in colorectal cancer, likely due to coregulation of Myc and Jun at the PIII upstream regulatory site.

### Acute Fbw7 loss in neural stem cells recapitulate findings from Hct116 cells

Because the Fbw7 mutations in the Hct116 cell panel were stably engineered into a transformed cell line, we examined the generalizability of these results by determining how acute Fbw7 deletion in a non-transformed cell line impacts RNA expression and Jun occupancy. We studied neural stem cells (NSCs) as a cell type with characterized Fbw7-mediated Jun regulation (Hoeck et al., 2010) and used a high efficiency CRISPR/nucleofection protocol to inactivate Fbw7 in U5 NSCs without the need for selection (Figure 6 – figure supplement 1A) (Hoellerbauer, Kufeld, & Paddison, 2020; Hoellerbauer, Kufeld, Arora, et al., 2020). Analogous to the Hct116 cell panel, ∼9% of protein-coding genes were differentially expressed in Fbw7^-/-^ cells compared with WT-U5 NSCs (Figure 6A). GO term analysis on the differentially expressed genes again revealed several enriched categories, some of which were shared with Hct116 cells (Figure 6B). Notably, “MHC class II” was one of the highly enriched GO terms in Fbw7^-/-^ U5 NSCs.

**Figure 6.**
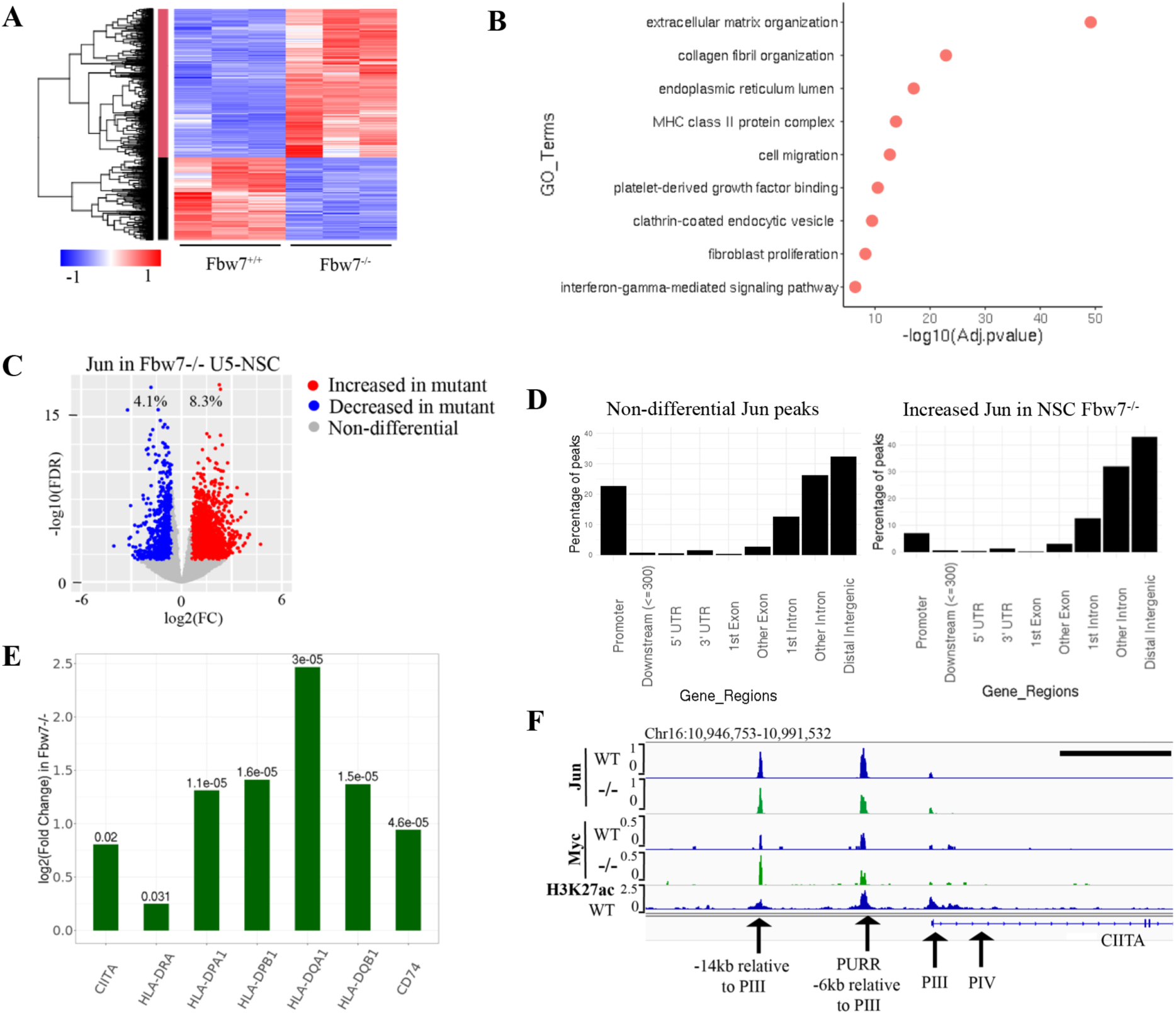
Transcriptional consequences of loss of Fbw7 in neural stem cells. (A) Clustering analysis separates differentially expressed protein-coding genes in NSCs into two groups. Heatmap shows the intensity of expression of each gene (y axis) for three replicates per cell type (x axis). (B) Gene Ontology terms enriched in differentially expressed genes in Fbw7^-/-^ NSCs. (C) Sites with increased (red) and decreased (blue) Jun in Fbw7^-/-^ NSCs compared to WT. (D) Non-differential and differential Jun peaks located within each gene feature. (E) Fold change of CIITA and MHC Class II genes in Fbw7^-/-^ NSCs compared to WT. FDR values are given at the top of each bar. n = 3 (F) Genome browser view of Myc, Jun, and H3K27ac occupancy on CIITA regulatory regions in WT and Fbw7^-/-^ NSCs. Black scale bar = 8 kb. See Figure 6 – figure supplement 1-3, Figure 6 – source data 1-5

We mapped Jun genomic occupancy in WT and Fbw7^-/-^ NSCs and found results that mirrored those in the Hct116 cell panel (Figure 6 – figure supplement 1B): (1) only a subset of the Jun binding sites displayed differential occupancy after Fbw7 inactivation (8.3% increased and 4.1% decreased) sites, and (2) most of the differentially regulated sites occurred in introns and intergenic regions (p value < 0.0001, Fisher test) (Figure 6C, 6D, Figure 6 – figure supplement 2). Thus, while the specific loci impacted by Fbw7 loss in the Hct116 cells NSCs differed, the scope of Fbw7’s impact on Jun was quite similar in both contexts. One striking similarity between the two systems was the regulation of CIITA and MHC class II expression (Figure 6E). Jun and Myc were both bound at regulatory regions upstream of CIITA in NSCs while Myc was differentially increased in Fbw7^-/-^ NSCs (Figure 6F). Unlike Hct116 cells, NSC WT cells express a basal level of CIITA, which may reflect constitutive (Fbw7-independent) Jun occupancy upstream of CIITA (Figure 6 – figure supplement 3).

## Discussion

Our primary goal was to understand the global transcriptional consequences of oncogenic Fbw7 mutations. Fbw7 loss affected ∼10% of all expressed genes, which could reflect Fbw7 substrates that are either sequence-specific TFs or global transcriptional regulators, such as the Mediator complex (M. A. Davis et al., 2013). However, only small subsets of the mapped H3K27ac and Jun/Myc binding sites were affected by the Fbw7 status. What might account for this specificity? One factor appears to be where the Fbw7-dependent loci are located, since most differential sites fell within distal regulatory elements. By targeting an enhancer rather than individual promoters, Fbw7 might cooperatively regulate multiple genes via a single regulatory region. TF phosphorylation may also be restricted to just the differential sites, thereby limiting the loci that can be targeted by Fbw7. If so, we might expect to find Fbw7 bound to these sites, as supported by our data implicating TFs in Fbw7 recruitment to chromatin (Figure 3 – figure supplement 1). However, we have not yet been able to map Fbw7 to specific chromatin sites. Despite the many differences between the Hct116 cell panel and NSCs (e.g., cell type, acute versus chronic Fbw7 loss, transformed versus non-transformed cells), Fbw7 loss had remarkably similar consequences, suggesting that these features reflect fundamental properties of Fbw7 function.

Although only a minority of Jun and Myc binding sites were differentially regulated by Fbw7, there was substantial overlap and remarkable co-regulation: in every case where they overlapped, both Jun and Myc occupancy were coordinately either increased or decreased by Fbw7 loss. We speculate that at coregulated sites such as CIITA, Fbw7 coordinately regulates transcription through the concerted targeting of both Jun and Myc. The expected outcome of Fbw7 loss is substrate accumulation, and most differential sites had increased occupancy in Fbw7 mutant cells. However, we also found differential sites with decreased Jun or Myc occupancy and correspondingly less mRNA expression in Fbw7 mutant cells. The mechanisms through which Fbw7 loss decreases TF occupancy remain to be elucidated. While Myc ubiquitylation regulates its role in transcriptional elongation (Jaenicke et al., 2016), our data do not presently provide insights into this potential aspect of Myc ubiquitylation by Fbw7.

Fbw7^R/+^ mutations may specifically stabilize those substrates that require a fully functional Fbw7 dimer (Welcker et al., 2013). One example may be Myc, which is partially stabilized by Fbw7^R /+^ in T-cells (King et al., 2013). Because excess Myc protein induces apoptosis, intermediate Myc stabilization caused by Fbw7^R/+^ may be an example of the “just enough” model (H. Davis & Tomlinson, 2012). Indeed, our finding that many of the deregulated genes and loci shared between the Fbw7^-/-^ and Fbw7^R/+^ mutant cell lines exhibited intermediate deregulation in the Fbw7^R/+^ cells supports the notion of “just enough” Fbw7^R/+^ transcriptional outcomes. However, we also found differential transcripts and loci that were more severely impacted by Fbw7^R/+^ than by Fbw7^-/-^, suggesting that heterozygous Fbw7 missense mutations also have unique functional outcomes (Figure 4B).

Our inability to link most differential sites to specific genes limited the identification of biologic pathways associated with Fbw7 loss. One important exception, however, is MHC Class II gene expression. Constitutive CIITA expression is normally confined to antigen presenting cells and it was striking to find CIITA and MHC Class II genes expressed in Fbw7 mutated colon cancer cells. Abnormal CIITA and MHC Class II expression has been observed in tumors, including colorectal cancers (Axelrod et al., 2019; Sconocchia et al., 2014). For example, aberrant CIITA expression in melanomas results from the activation of both IFNγ-inducible and constitutive CIITA promoters (van der Stoep et al., 2007). Moreover, deletion of the AP-1 site that we found differentially occupied by Jun in Fbw7 mutant cells was found to compromise CIITA expression in melanoma cells (van der Stoep et al., 2007). In sum, these data support a model in which Fbw7 loss indirectly augments CIITA expression through increased Jun and Myc occupancy at these regulatory regions.

What are the implications of Fbw7-dependent CIITA expression in colorectal cancers? Tumor cell specific MHC Class II expression is generally associated with favorable prognosis, which may reflect increased tumor immunogenicity conferred by MHC Class II expression. Intriguingly, *FBXW7* mutations and distant metastasis almost never co-occur in colorectal cancers (Muzny et al., 2012). Fbw7 loss may thus confer better prognosis, perhaps due to Fbw7-dependent MHC class II upregulation. Accordingly, we found increased CIITA expression in TCGA colorectal tumors and CCLE colorectal cancer cell lines with *FBXW7* mutations. Previously, we used machine learning to develop gene expression signatures that predicted *FBXW7* mutations in TCGA tumors. While we focused on a metabolic gene signature in the study, we also found that a signature comprised of MHC Class II and other genes associated with immune response was also highly predictive of *FBXW7* mutations in colorectal cancers (R. J. Davis et al., 2018 Supplemental Dataset S02). While these TCGA analyses are correlative, in light of our finding that Fbw7 loss induces CIITA expression in Hct116 cells and NSCs, we speculate that Fbw7 mutations in colorectal cancers lead to increased CIITA expression, increased immunogenicity, and better prognosis.

Others have also found associations between Fbw7 status and immune responses in cancer. A recent study found that an Fbw7^R/+^ mutation conferred resistance to PD-1 blockade through impaired dsRNA sensing and IFNγ signaling in a metastatic melanoma and a murine melanoma model (Gstalder et al., 2020). Fbw7 loss also correlated with decreased IFNγ signaling in TCGA cancers. In this case, Fbw7^R/+^ decreased MHC Class I but not MHC Class II expression and caused a more aggressive phenotype associated with decreased immunogenicity. These discrepancies with our study may relate to the different model systems and tumor types. In mice, Fbw7 also regulates the tumor microenvironment through a non-tumor-cell-autonomous manner involving expression of the CCL2 chemokine (Yumimoto et al., 2015). Further studies are needed to fully appreciate the pathologic and therapeutic implications of Fbw7-related tumor immunogenicity and immune responses to cancer.

Overall, these data establish a framework for understanding how mutations in Fbw7 can exert context-specific deregulatory effects and how Fbw7 substrates can act synergistically to drive tumor progression.

## Materials and Methods

### RNA-Seq: RNA isolation, library preparation, sequencing and data analysis

RNA was isolated using the Qiagen RNeasy Mini Kit (Cat# 74104) following the manufacturer’s instructions. Three replicates per cell type were included and for each replicate cells were harvested from separate cultured plates. RNA quality and integrity were determined (A260/280 1.8 – 2.1, A260/230 > 1.7, RIN ≥ 9). Libraries were prepared by the Fred Hutch Genomics Center using the TruSeq RNA Samples Prep Kit v2 (Illumina Inc., San Diego CA, USA). Sequencing was performed on a Illumina HiSeq 2500 with 50 bp paired-end reads (PE50). RNA-Seq for U5-NSCs was an exception. Knockouts were generated separately on two different days and cells from separate nucleofection reactions were used as the three replicates, hence biological replicates. Libraries were prepared using TruSeq Stranded mRNA and sequencing was performed using an Illumina NovaSeq 6000 employing a paired-end 50 base read length (PE50).

Fastq files were filtered to exclude reads that didn’t pass Illumina’s base call quality threshold. STAR v2.7.1 (Dobin et al., 2013) with 2-pass mapping was used to align paired-end reads to human genome build hg19 and GENCODE gene annotation v31lift37 (https://www.gencodegenes.org/human/). FastQC 0.11.8 (https://www.bioinformatics.babraham.ac.uk/projects/fastqc/) and RSeQC 3.0.0 (Wang et al., 2012) were used for QC including insert fragment size, read quality, read duplication rates, gene body coverage and read distribution in different genomic regions. FeatureCounts (Liao et al., 2014) in Subread 1.6.5 was used to quantify gene-level expression. For stranded libraries, only coding strand derived reads were counted. Bioconductor package edgeR 3.26.8 (Robinson et al., 2009) was used to detect differential gene expression between conditions. Genes with low expression were excluded by requiring at least one count per million in at least N samples (N is equal to one less than the number of samples in the smallest group). The filtered expression matrix was normalized by TMM method (Robinson & Oshlack, 2010) and subject to significance testing using generalized linear model and quasi-likelihood method. Genes were deemed differentially expressed if absolute fold changes were above 1.5 and FDRs were less than 0.05.

### Cleavage Under Target and Release Using Nuclease (CUT&RUN)

Manual or automated CUT&RUN were performed as previously described (Janssens et al., 2018; Skene et al., 2018; Skene & Henikoff, 2017). Briefly, cells were harvested using Accutase, counted and washed twice with Wash Buffer (20mM HEPES pH 7.5, 150 mM NaCl, 0.5mM Spermidine and one Roche Complete EDTA free protein inhibitor tablet per 50 mL). Cells were bound to Concanavalin A-coated magnetic beads (20uL per one million cells). Then cells were permeabilized with Dig Wash buffer (Wash Buffer with 0.05% Digitonin) while being incubated with primary antibody overnight at 4 ^ο^C. Cell-bead mixture was washed twice with Dig-Wash buffer and incubated with Protein A-MNase (pA-MN) for 1 hour at 4 ^ο^C. After washing the mix with Dig Wash buffer twice, cells were placed on an ice-cold block and incubated with 2 mM CaCl_2_ in Dig Wash buffer to activate pA-MN digestion. After the specific digestion period the reaction was inhibited with 2X Stop Buffer (340 mM NaCl, 20mM EDTA, 4mM EGTA, 0.05% Digitonin, 0.05% mg/ml glycogen, 5 ug/mL RNase, 2pg/mL heterologous spike-in DNA). The samples were incubated at 37 ^ο^C for 30 min to release the digested DNA fragments into the supernatant. The supernatant was collected and libraries were prepared as previously explained (Janssens et al., 2018). Paired-end 25 base read length (PE25) sequencing was performed using an Illumina HiSeq 2500 platform at Fred Hutch Genomics Shared Resources.

<u>Deviations from the above described protocol:</u>

1. Automated CUT&RUN: Manual preparation included harvesting cells, counting, washing, permeabilizing and antibody addition. After cells were incubated with the antibody at 4 ^ο^C overnight, next day the samples were submitted for automated CUT&RUN which was performed by the Genomics and Bioinformatics Center at Fred Hutch on a BioMek platform.
2. Used nuclei instead of cells: H3K4me1 and H3K4me2 were mapped using the CUT&RUN protocol as previously described using isolated nuclei (Skene & Henikoff, 2017).

A summary of all CUT&RUN samples with conditions and method used can be found at Additional source data 1.

### CUT&RUN data analysis

Basic analysis: Sequencing reads were aligned to hg19 using Bowtie2: bowtie2 --end-to-end -- very-sensitive --no-overlap --no-dovetail --no-unal --no-mixed --no-discordant -q -I 10 -X 700 -x path/to/Bowtie2/indices -1 read1.fastq.gz -2 read2.fastq.gz

CPM normalized bigwig files were generated using bedtools genomecov.

Peaks were called using MACS2. Peak calling was performed for each target with and without the IgG control.

Narrow peaks with IgG control: macs2 callpeak --name TARGET --treatment path/to/TARGET/hg19.bam --control path/to/IgG/hg19.bam --format BAMPE --gsize hs --keep-dup all -q 0.05

Narrow peaks without IgG control: macs2 callpeak --name TARGET --treatment path/to/TARGET/hg19.bam --format BAMPE --gsize hs --keep-dup all -q 0.05

IgG-controlled peaks that overlap with no-control peaks were retained for further analyses. For each TF/histone mark mapped in each genotype, peaks from three replicates were considered to make a final peak-set to use for downstream analysis.

Differential binding analysis: Merged peak set for each target was used for the analysis. FeatureCounts (Liao et al., 2014) in Subread 1.6.5 was used to count reads mapped to merged peaks in each sample. Bioconductor package edgeR 3.26.8 (Robinson et al., 2009) was used to detect differential peaks between conditions. Peaks with low read numbers were excluded using edgeR function filterByExpr with min.count = 10 and min.total.count = 15. The filtered count matrix was normalized by TMM method (Robinson & Oshlack, 2010) and subjected to significance testing using generalized linear model and quasi-likelihood method. Peaks were deemed differentially bound if absolute fold changes were above 1.5 and FDRs were less than 0.1 for H3K27ac and Jun data, and FDR 0.05 for Myc data. Differential sites for H3K27ac, Jun and Myc are provided as source data.

### Other data processing, analysis and visualization

1. Correlation between RNA-Seq and the distribution of histone marks around Transcriptional Start Sites (TSSs).

A reference list of hg19 genes was downloaded from the UCSC Table Browser. Genes were oriented according to the directionality of gene transcription and specified a 2 kb window around TSSs. Genes that have an overlapping TSS within the 2 kb window and mitochondrial genes were removed, creating a list of 22,222 TSSs. The gene list was sorted in descending order of their RNA-Seq FPKM values. CUT&RUN H3K27ac and H3K27me3 signal (merged from three replicates) were mapped on to the ordered genomic sites. The coverage of histone marks was quantified using bedtools coverage and converted to FPKM values. Correlation between RNA-Seq and histone mark FPKM values was calculated using R cor.test function (method=spearman).

2. Correlation matrices were generated using deepTools (Ramírez et al., 2016).
3. Gene set enrichment analysis (Gene Ontology terms) was done using the Enrichr web-based tool (Kuleshov et al., 2016).
4. Motif identification. For all motif analysis we used the The MEME Suite (Bailey et al., 2009). We used bedtools getfasta to generate FASTA files for genomic sites of interest (Quinlan & Hall, 2010). For motif discovery analysis we submitted the center 100 bp sequence of peaks to MEME-ChIP. MEME-ChIP was used with default parameters in Classic mode. HOCOMOCO Human (v11 FULL) motif database was used. We used the position-weight matrix (PWM) of the motif discovered by MEME-ChIP as the input for FIMO, to quantify the abundance of the motif. We used FIMO with a threshold value of p.value ≤ 0.01 to capture all motif configurations and then filtered the output to select only the motifs with the highest FIMO motif scores (higher the score, similar to the input motif). For differential motif analysis, we used MEME-ChIP in Differential Enrichment mode with default parameters.
5. Annotations. To assign gene regions where peaks are located, we used ChIPseeker, an R/Bioconductor package (Yu et al., 2015). We used the nearest gene method to assign a peak to a gene using the bedtools closest tool (Quinlan & Hall, 2010). Gencode Human Release 31 (GRCh37) Comprehensive gene annotation list was used to generate a list of genes with full gene coordinates which was used to annotate peaks to the nearest gene.
6. Data Visualization. Plots were generated using R (https://www.r-project.org) (R Core Team, 2020). Heatmaps were generated using Deeptools (Ramírez et al., 2016).
7. Venn diagrams. Intersection between genomic sites were generated using Intervene Venn module (A. Khan & Mathelier, 2017).
8. Primary cancer and cell line data analysis. CIITA expression data from Fbw7 WT and mutated colon and rectal cancers were collected from the TCGA COADREAD database via UCSC Xena browser (Goldman et al., 2020) (Figure 5 – source data 2). CIITA expression in Fbw7 WT and mutated colorectal cancer cell lines were collected from the DepMap Portal (https://depmap.org/portal/) (Barretina et al., 2012). Statistical analysis was performed on GraphPad Prism. Unpaired t-test (two tailed) was used to determine statistical significance of CIITA differential expression of TCGA and CCLE data sets.
9. Bigwig files (three replicates merged) were viewed on Integrative Genome Viewer to show examples of CUT&RUN binding data as peaks. Schematic figures were created with BioRender.com.

### Antibodies

For CUT&RUN we used Rabbit anti-H3K27ac (1:100, Abcam Cat #ab45173), anti-H3K27me3 (1:100, Cell Signaling Technologies Cat#9733S), Rabbit c-Jun (1:25, Santa Cruz Cat #sc-1694), Rabbit anti-Myc (1:25, Cell Signaling Technologies D3N8F Cat #13987), Rabbit anti-H3K4me1 (1:100, Abcam Cat#ab8895), Rabbit anti-H3K4me2 (1:100, Cell Signaling Tech 9725) and (Rabbit normal IgG (1:50, Santa Cruz sc-2027). For western blots and immunoprecipitation we used anti-Fbw7 Bethyl A301-720A.

### Cell culture

Hct116 cells were grown in DMEM with 10% FBS and 5% PenStrep. For CUT&RUN and RNAseq experiments 2 x 10^6^ cells were plated per 10 cm dish two days prior to harvesting. Cells were harvested using Accutase. Human fetal tissue derived U5 NCSs were cultured in NeuroCult NS-A basal medium (Stem Cell Technologies) supplemented with N2 (made in-house 2x stock in Advanced DMEM/F-12 (Thermo Fisher Scientific)), B27 (Thermo Fisher Scientific), antibiotic-antimycotic (Thermo Fisher Scientific), glutamaz (Thermo Fisher Scientific), EGF and bFGF (Peprotech). Cells were cultured in Laminin coated plates. Accutase was used to harvest cells for experiments.

### Chromatin fractionation

Untreated and Bortezomib treated (0.5 uM for 10 hrs.) cells were harvested and counted. Cells were resuspended in CSK buffer (10 mM HEPES pH 6.8, 100mM NaCl, 1mM EGTA, 1mM EDTA, 2mM MgCl_2_, 300mM Sucrose, 0.1% Triton X-100 and Protease inhibitor - 50 ul per million cells) (Kim et al., 2008). Cells were allowed to lyse for 5 min on ice and centrifuged for 5 min at 4^0^C at 1500 g. The supernatant which is the soluble fraction (S) was removed to a new tube. The pellet was resuspended in 1 ml of CSK buffer, centrifuged for 5 min at 4^0^C at 1500 g. The supernatant was thoroughly removed. Next, NP40 buffer with protease inhibitor and 250 U/ml benzonase was added to the cell pellet (same volume as CSK buffer was used to lyse cells). Cells were incubated for 30 min on ice. This was the chromatin fraction (C). Both soluble and chromatin fractions were sonicated and centrifuged to remove debris (5 min at 4^0^C at maximum speed). Total protein in all chromatin fractions was quantified using the Bradford assay and samples were normalized to total protein content. Equal volumes of chromatin and soluble fractions from each sample were used to immunoprecipitate Fbw7. Chromatin fractionation of Fbw7 was determined by >3 independent experiments.

### Immunoprecipitations and Western blot analysis of Fbw7

Whole cell extracts (WCE) were made by lysing cells in 0.5% NP-40 buffer with protease inhibitor cocktail (made in-house). Then WCE were sonicated and spun to remove debris. To immunoprecipitate Fbw7 from whole or fractionated cell lysates anti-Fbw7 Bethyl A301-720A antibody and Protein A beads were added and incubated for at least 2hrs at 4^0^C (overnight recommended). Beads were then washed 3X with 1 ml NP40 lysis buffer. Eluted protein was electrophoresed on 8% polyacrylamide gels and transferred to PVDF which was blotted against Fbw7 using anti-Fbw7 Bethyl A301-720A (1:1000) and HRP conjugated anti-Rabbit secondary antibody (1:10,000). Membranes treated with ECL (made in-house) were visualized on a BioRad ChemiDoc imaging system.

### PCR amplification of CIITA

RNA was isolated from Hct116 and Raji cells using the Qiagen RNeasy Mini Kit (Cat # 74104). cDNA was prepared using the iScript Reverse Transcription Supermix (Cat # 1708841). CIITA PIII and PIV were amplified using specific primers (PIII : F – 5’GCTGGGATTCCTACACAATGC3’, R – 5’GGGTTCTGAGTAGAGCTCAATC3’ and PIV : F – 5’GGGAGCCCGGGGAACA3’, R – 5’GATGGTGTCTGTGTCGGGTT3’) at 60^0^C annealing temperature for 38 cycles (H. Chen et al., 2015). GAPDH was amplified as the control (25 cycles) using primers F – 5’GGTCGGAGTCAACGGATTTG3’ and R – 5’ATGAGCCCCAGCCTTCTCCAT3’. Platinum Taq DNA polymerase was used following the manufacturer’s instructions.

### Generation of U5-NSC homozygous Fbw7 knockouts

A previously described protocol to generate homozygous null mutations using CRISPR-Cas9 and nucleofection was followed (Hoellerbauer, Kufeld, & Paddison, 2020; Hoellerbauer, Kufeld, Arora, et al., 2020). Briefly the protocol is as follows:

CRISPR sgRNA were designed using Broad Institute’s GPP Web Portal. The output list of sgRNAs was manually curated to choose three sgRNAs targeting *FBXW7*. Exon 3, 4 and 9 in *FBXW7* were targeted by 5’AAGAGCGGACCTCAGAACCA3’, 5’CTGAGGTCCCCAAAAGTTGT3’, 5’ACATTAGTGGGACATACAGG3’ guides respectively. A control sgRNA was included 5’GTAGCGAACGTGTCCGGCGT3’. sgRNAs were purchased from Synthego.

#### Cas9:sgRNA RNP nucleofection

Reconstituted sgRNAs by adding 10uL of 1X TE Buffer 1.5nmol of dried sgRNA. A working stock of 30uM sgRNA was used henceforth. A working stock of Cas9 (10.17 pmol/ul) was made. To prepare RNP complexes, sgRNA, sNLS-SpCas9-sNLS (Aldevron) and SG Cell Line Nucleofector Solution (Lonza) were mixed in 1.87 uL, 1.84 uL and 18.29 uL respectively to make a 22 uL final volume. The mixture was incubated at room temperature for 15 minutes to allow RNP complexes to form. To nucleofect, 0.13 x 10^6^ cells were harvested. The cells were washed with PBS and resuspended with RNPs. (We were able to successfully nucleofect up to 0.85 x 10^6^ cells with the same volume of RNPs.) Cells were electroporated using the Amaxa 4D Nucleofector X unit and program EN-138. Nucleofected cells were plated in pre-warmed media.

#### CRISPR editing efficiency analysis

Extraction of genomic DNA, PCR amplification of target site and efficiency analysis was done as previously described (Hoellerbauer, Kufeld, & Paddison, 2020; Hoellerbauer, Kufeld, Arora, et al., 2020). The primer pairs used to amplify CRISPR target sites in Exon 3: 5’TCATCACACACTGTTCTTCTGGA3’ and 5’TGTCTACCCTAGAACAGCTGT3’, Exon 4: 5’TGTGTACCTGTGATCTCTGGG3’ and 5’CACCTTGCTGTGCAACCATC3’, Exon 9: 5’ACTGCTTTCATGTCGTGTTTCC3’ and 5’AGGAAGCTGACAACACTAGCA3’. We found that the pool of three sgRNA was the most successful at deleting *FBXW7*. It was confirmed by blotting for immunoprecipitated Fbw7 in each nucleofected sample (Figure 6 – figure supplement 1).

## Data Availability

All data generated and used in this manuscript are deposited in GEO: GSE184041

Scripts available at https://github.com/hnthirima

The results shown here are in part based upon data generated by the TCGA Research Network: https://www.cancer.gov/tcga

## Source data files (with title and legend)

**Figure 1 - source data 1: Differential expression analysis of Hct116 RNA-Seq**. This excel file contains the differential analysis output of Hct116 RNA-Seq data from WT, Fbw7^-/-^ (Del) and Fbw7^R/+^ (R). DE = 0, not differentially expressed in the mutant compared to WT; DE = 1, differentially expressed in the mutant compared to WT.

**Figure 1 - source data 2: Hierarchical cluster output file.** This excel file includes genes that belong to each cluster in the hierarchical cluster analysis.

**Figure 1- source data 3: Enrichr output for Hct116 differentially expressed genes.** This excel file includes the GO Terms enriched in differential genes unique to Fbw7^-/-^ (Cluster 1 and 2) and, common to Fbw7^-/-^ and Fbw7^R/+^ (Cluster 3 and 4). Genes in each GO Term are listed along with P-value and Adjusted P-value.

**Figure 2 – source data 1**: **H3K27ac differential sites**. This excel file includes lists of peaks with increased and decreased H3K27ac signal in Hct116 Fbw7^-/-^ and Fbw7^R/+^ relative to WT. Fold change and FDR listed for each peak.

**Figure 2 – source data 2: Summary of CUT&RUN differential sites.** This excel file includes a summary (total number of differential sites, percentage, number of annotated genes) of H3K27ac, Jun and Myc differential sites in Hct116 cells and Jun differential sites in U5-NSCs.

**Figure 3 – source data 1: Original western blots for** **Figure 3A** **and Figure 3 – figure supplement 1.**

**Figure 3 – source data 2: Jun and Myc differential sites in Hct116 cells.** This excel file includes lists of peaks with increased and decreased Jun and Myc signal in Hct116 Fbw7^-/-^ and Fbw7^R/+^ relative to WT. Fold change and FDR listed for each peak.

**Figure 5 – source data 1: Original gels for** **Figure 5C**.

**Figure 5 – source data 2: TCGA COADREAD data used for** **Figure 5D**. This excel file includes the CIITA expression counts for WT and Fbw7-mutant Colorectal tumors.

**Figure 5 – source data 3: Colorectal cancer cell line data from DepMap used for** **Figure 5E**.

**Figure 6 – source data 1: Confirming loss of Fbw7 in U5-NSC Fbw7^-/-^ cells.** This folder contains the original western blots for Figure 6 – figure supplement 1A. Western blots that confirm the loss of Fbw7 in two other separately performed nucleofection reactions are also included.

**Figure 6 – source data 2: Differential expression analysis of U5-NSC RNA-Seq.** This excel file contains the differential analysis output of U5-NSC RNA-Seq data from control (Ctrl23, Ctrl4.1, Ctrl4.2) and Fbw7^-/-^ (Fb23, Fb4.1, Fb4.2) cells. DE = 0, not differentially expressed in the mutant compared to WT; DE = 1, differentially expressed in the mutant compared to WT.

**Figure 6 – source data 3: Enrichr output for U5-NSC differentially expressed genes.** Genes in each GO Term are listed along with P-value and Adjusted P-value.

**Figure 6 – source data 4: Jun differential sites in U5-NSCs.** (U5F= Fbw7^-/-^ and U5W = WT)

**Figure 6 – source data 5: Original gels for Figure 6 – figure supplement 3.**

**Additional source data 1: Summary of all CUT&RUN experiments**. Experimental conditions of all CUT&RUN experiments included in the study.

## Acknowledgments

This research was supported by the following grants: NCI/NIH T32 CA080416 (H.N.T.), NCI/NIH R01 CA215647 (B.E.C.), R01 HG010492 (S.H.), R01NS119650 and R01 CA190957 (P.J.P) and the Genomics & Bioinformatics Shared Resource of the Fred Hutch/University of Washington Cancer Consortium (P30 CA015704).

We thank Jeff Delrow, Matthew Fitzgibbon, Alyssa Dawson and Philip Corrin in the Genomics Shared Resources at Fred Hutchinson Cancer Research Center for support with experimental design, helpful discussions, library preparation and sequencing. We thank Markus Welcker and Ahmed Diab for helpful discussions and critical reading of the manuscript. We thank past and present Henikoff lab members including Jorja Henikoff, Christine Codomo, Michael Meers, Jay Sarthy, Terri Bryson and Peter Skene for helpful discussions regarding data analysis, reagents, library preparation and sequencing. We also thank Pia Hoellerbauer in Paddison lab for assistance with neural stem cell culturing and knockout generation.

## Supplemental Figures

**Figure 1 – figure supplement 1.**
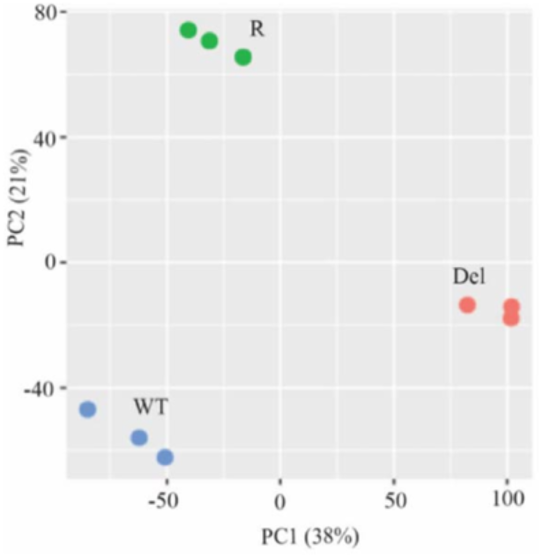
Principal Component Analysis (PCA) of RNA-Seq from Hct116 cells. Hct116 wild-type (WT), Fbw7^-/-^ (Del) and Fbw7^R/+^ (R) samples separate by genotype. Replicates from each condition are clustered together.

**Figure 2 – figure supplement 1.**
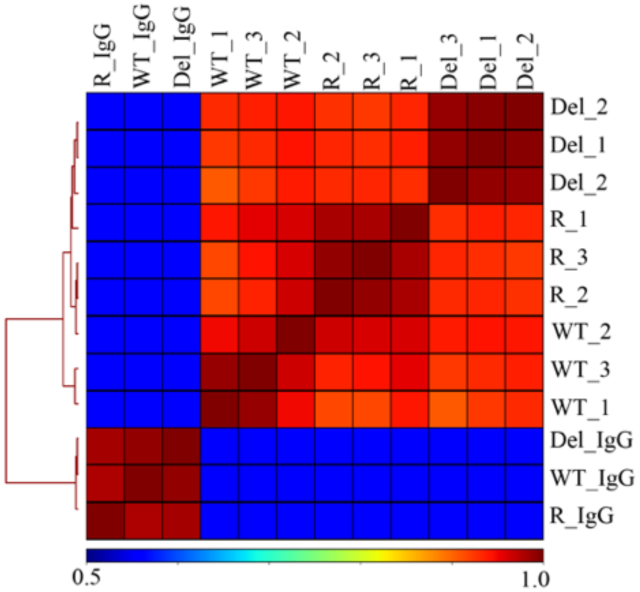
Hierarchically clustered correlation matrix of H3K27ac CUT&RUN profiles in Hct116 cells. Correlation matrix of three replicates from Hct116 WT, Fbw7^-/-^ (Del) and Fbw7^R/+^ (R) cells. IgG negative control for each cell type included. Peaks from the three cell types were merged to create a final peak-set to perform the correlation analysis.

**Figure 2 – figure supplement 2.**
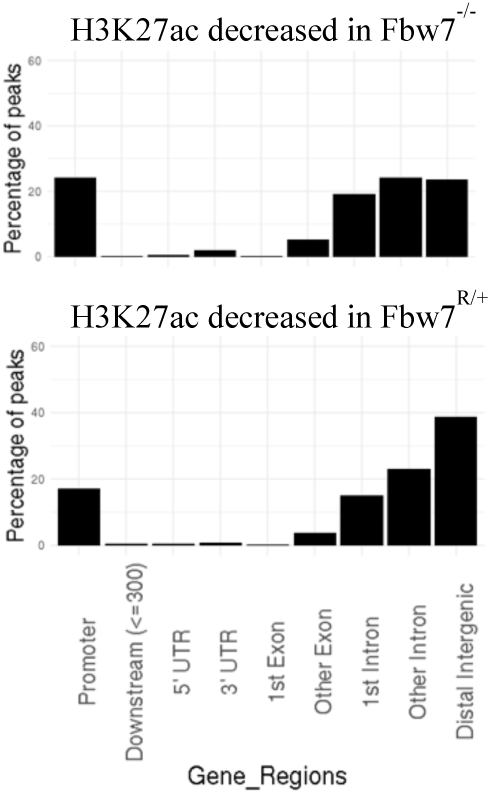
Percentage of peaks with decreased H3K27ac signal located within different gene features. Compared to non-differential H3K27ac peaks, differential H3K27ac peaks enrich mostly within introns and intergenic regions (p value < 0.0001, Fisher test).

**Figure 2 – figure supplement 3.**
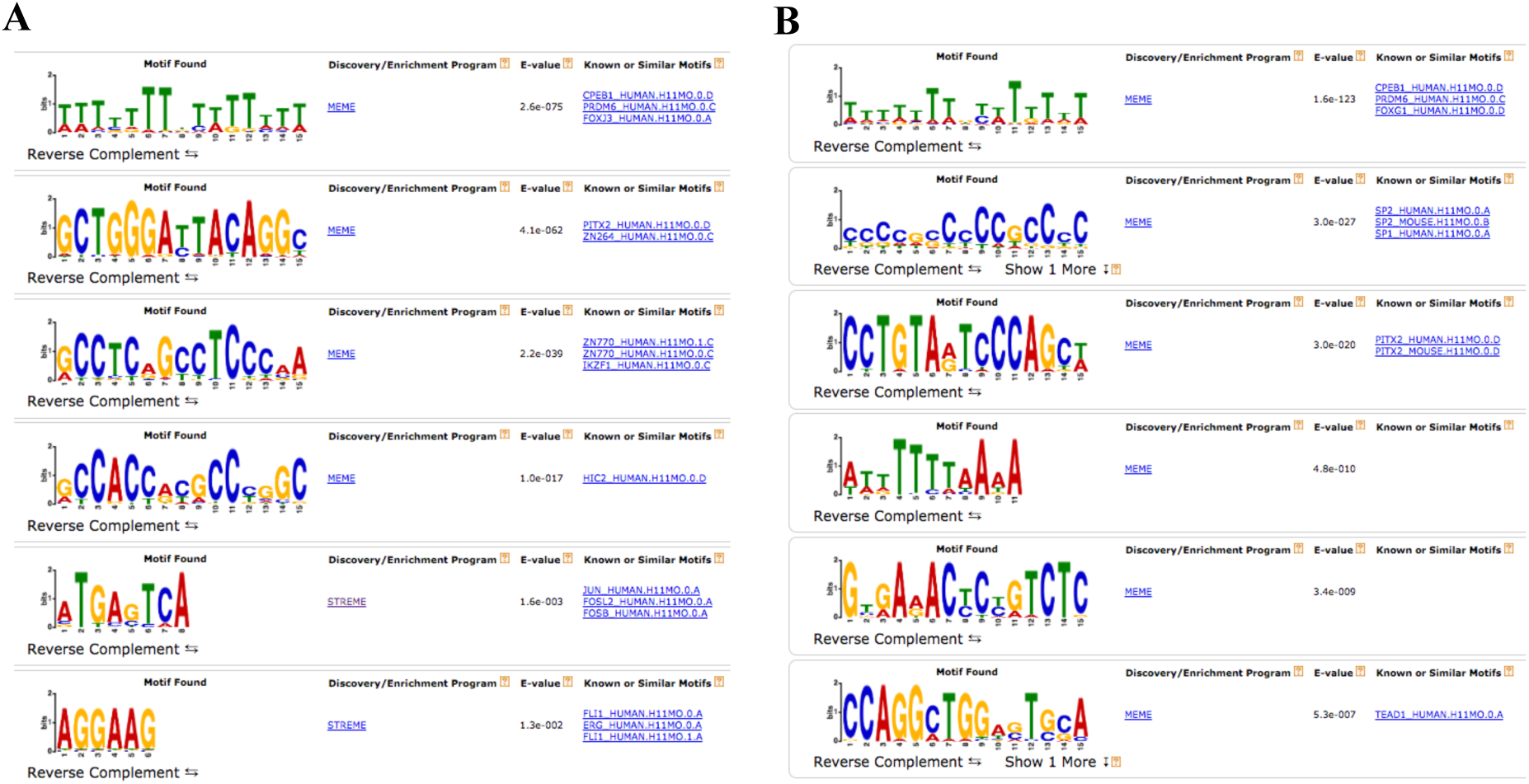
Complete output of the MEME-ChIP analysis on H3K27ac differential sites. (A) MEME-ChIP analysis on the sequences of H3K27ac increased sites in Fbw7^-/-^ cells. (B) MEME-ChIP analysis on the sequences of non-differential H3K27ac sites in Fbw7^-/-^ cells (negative control, 1409 sites). *FIMO analysis revealed that AP-1 motif was enriched in approximately 30-35% of H3K27ac sites that were decreased in Fbw7^-/-^ (p value ≤ 1.8e-5), increased in Fbw7^R/+^ (30.2% p value = 1.8e-5), decreased in Fbw7^R/+^ (35% p value ≤ 1.5e-5), however only 17% in non-differential sites (1409 sites) (p value ≤ 1.8e-5).

**Figure 3 – figure supplement 1.**
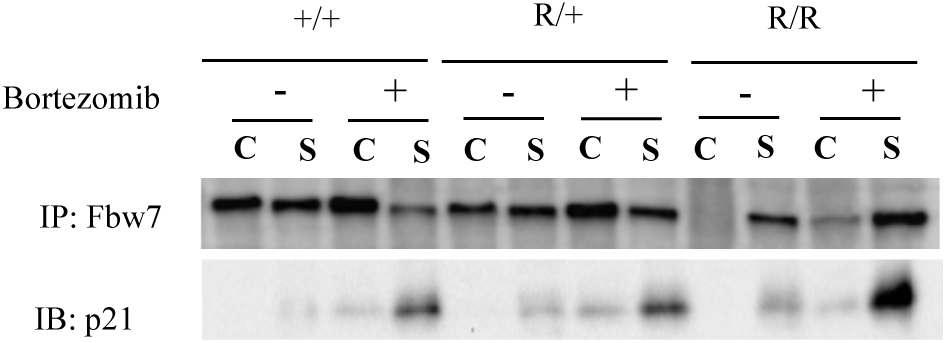
Fbw7 abundance in chromatin (C) and soluble (S) fractions from Hct116 WT, Fbw7^R/+^ and Fbw7^R/R^ cells treated with and without Bortezomib.

**Figure 3 – figure supplement 2.**
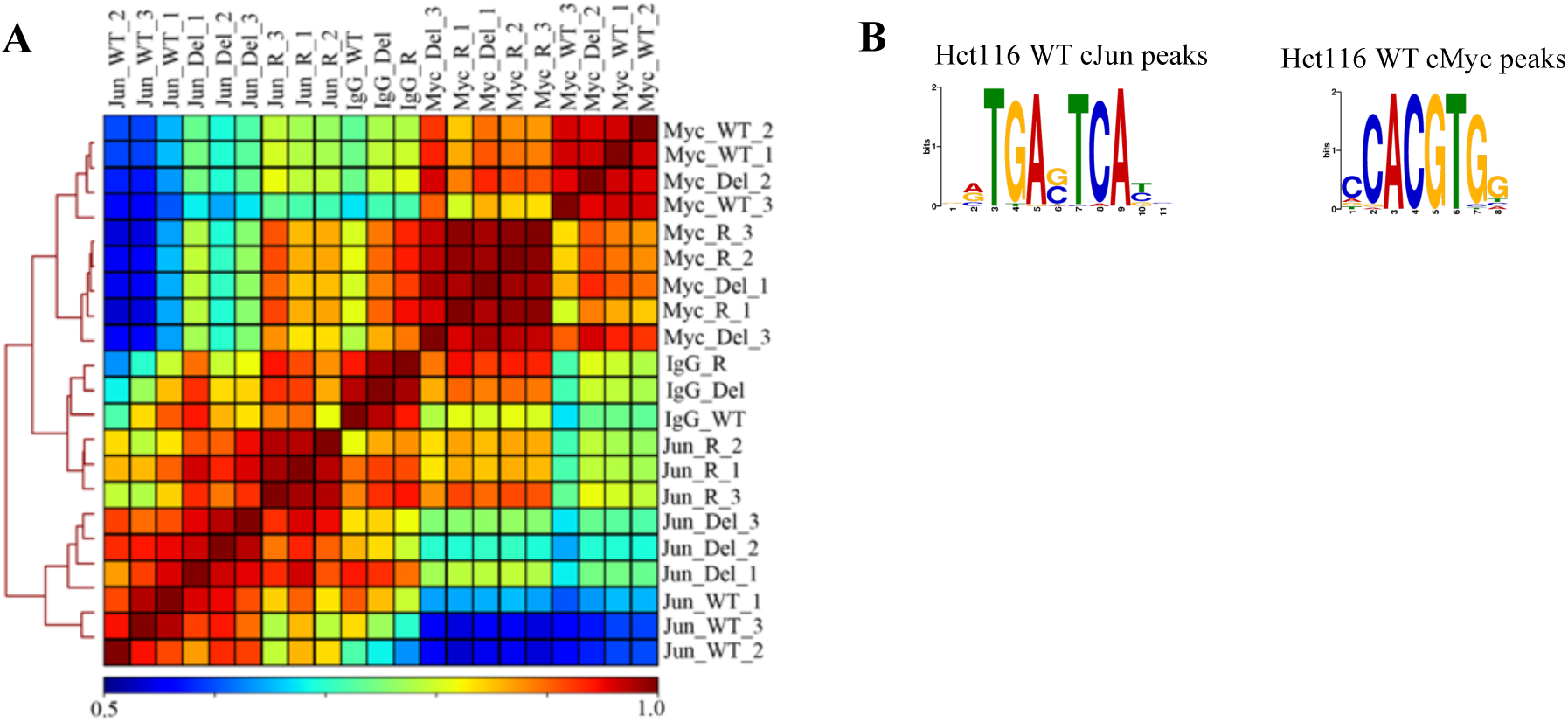
Validation of Jun and Myc CUT&RUN profiles. (A) Hierarchically clustered correlation matrix of Jun and Myc signal mapped in Hct116 WT, Fbw7^-/-^ (Del) and Fbw7^R/+^ (R) cells. IgG negative control for each cell type included. Peaks from the three cell types were merged to create a final peak-set. (B) Sequence logo of the AP-1 motif enriched in the center 100bp sequence of Jun peaks in Hct116 WT (E value 1.3e-53) and sequence logo of E-box motif enriched in the center 100bp sequence of Myc peaks in Hct116 WT (1.7e-4). AP-1 motif was input to FIMO to scan for the motif in full sequence of 25,527 Jun peaks in Hct116 WT. FIMO output showed that motifs with score between15.73 to 12.11 occurred 26,547 times (p value ≤ 6.46E-05). E-box motif was input to FIMO to scan for the motif in full sequence of 24,111 Myc peaks in Hct116 WT. FIMO output showed that motifs with score between 15.32 – 9.24 occurred 9343 times (p value ≤ 0.00024). Motif score range was determined by the exact similarity to TGAG/CTCA (AP-1 motif) or CACGTG (E box).

**Figure 3 – figure supplement 3.**
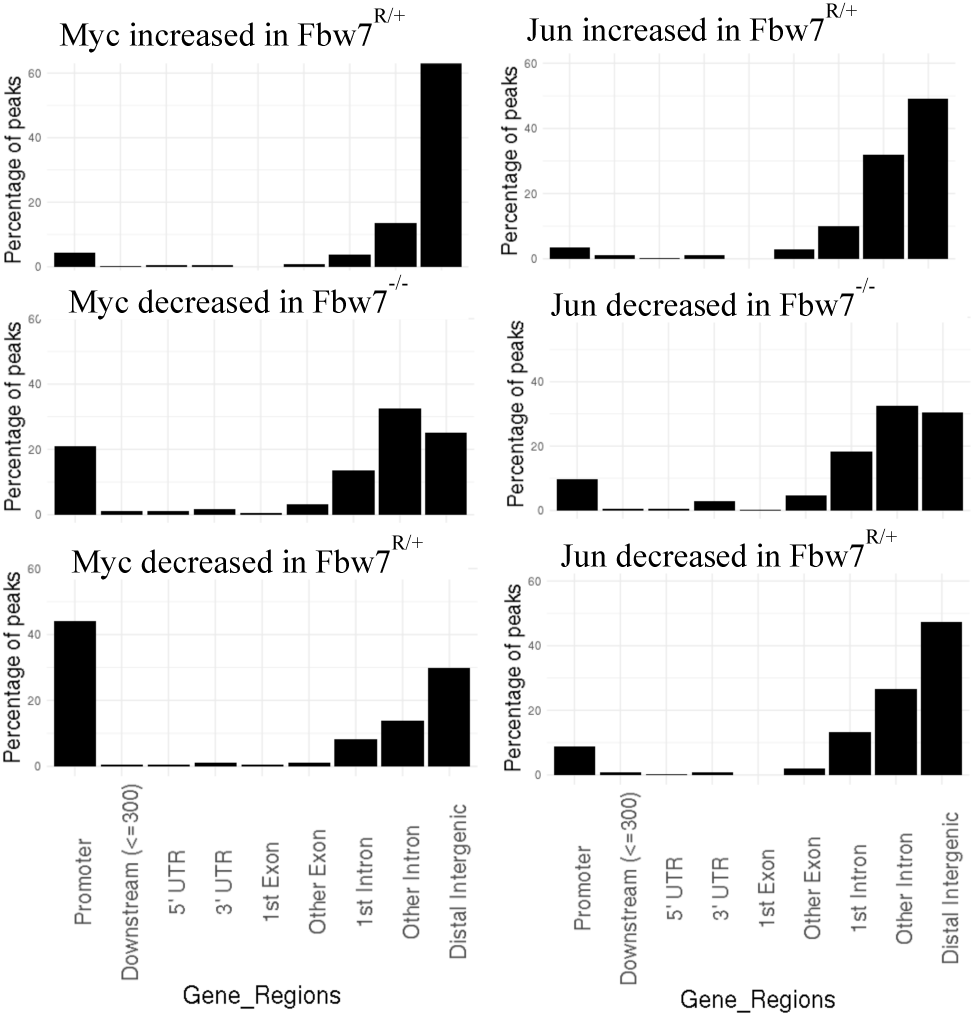
Percentage of Myc and Jun peaks located at different gene features.

**Figure 3 – figure supplement 4.**
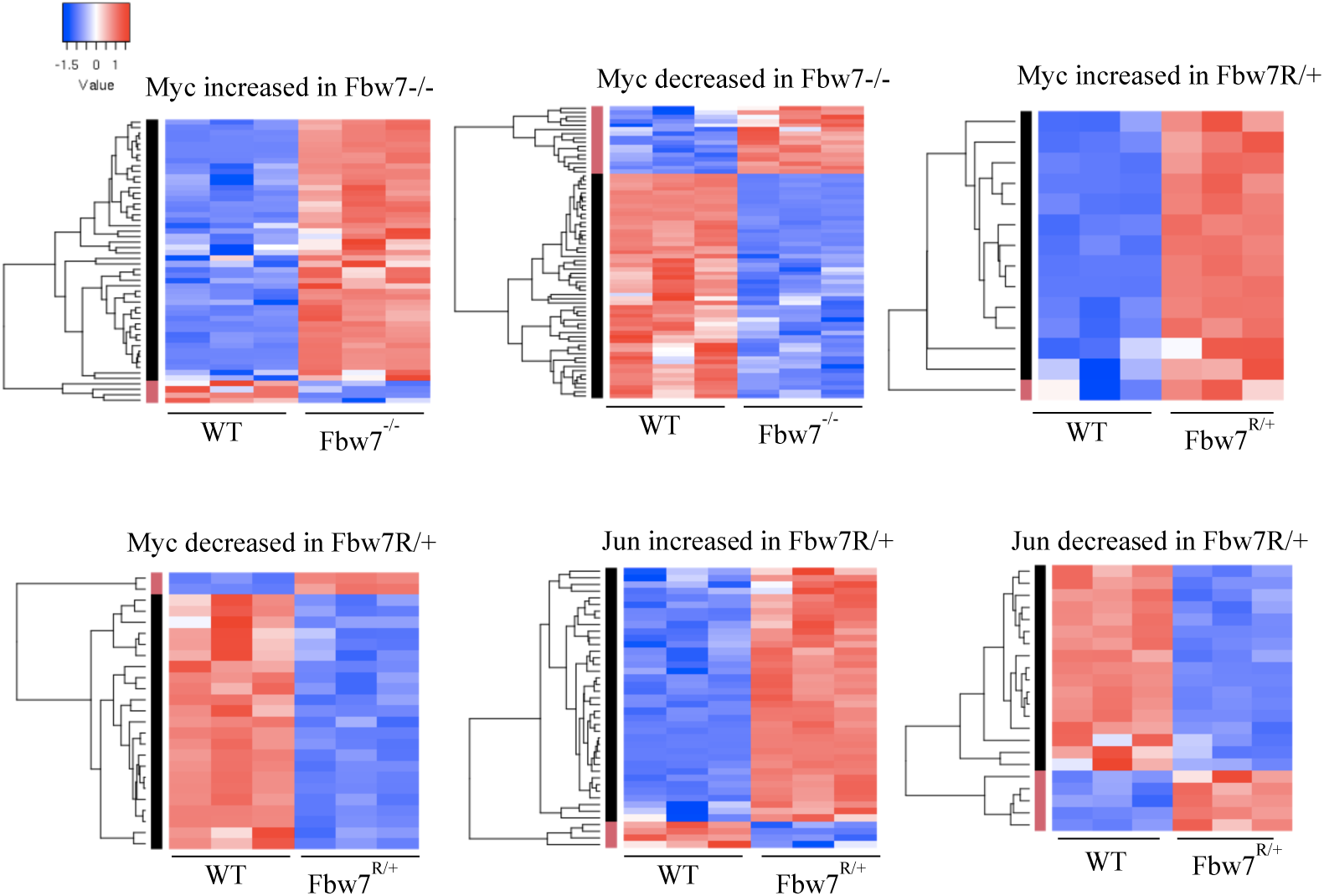
Transcription of genes with differential promoter proximal Myc and Jun occupancy in Fbw7 mutant cells. Hierarchically clustered genes showing the transcription of genes that have increased or decreased Myc and Jun occupancy at gene proximal sites in Fbw7^-/-^ and Fbw7^R/+^ cells.

**Figure 4 – figure supplement 1.**
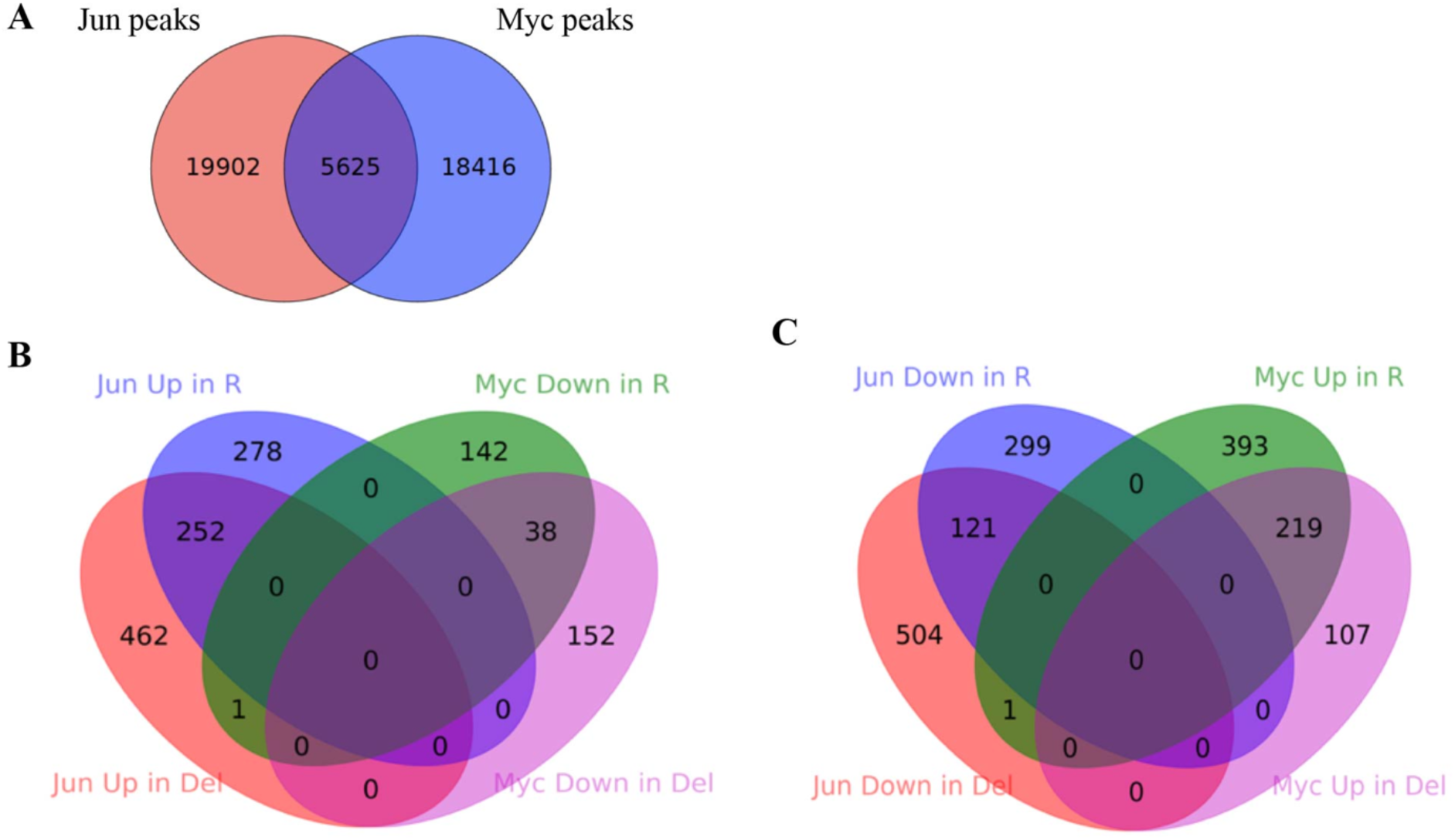
Comparison between Jun and Myc peaks in Hct116 cells. (A) The overlap between Jun and Myc peaks in Hct116 WT cells. (p value < 0.0001, Fisher Test) (B) The overlap between peaks with increased Jun occupancy in Fbw7^-/-^ and Fbw7^R/+^ cells, and decreased Myc occupancy in Fbw7^-/-^ and Fbw7^R/+^ cells (C) The overlap between decreased Jun occupancy in Fbw7^-/-^ and Fbw7^R/+^ cells, and increased Myc occupancy in Fbw7^-/-^ and Fbw7^R/+^ cells.

**Figure 5 – figure supplement 1.**
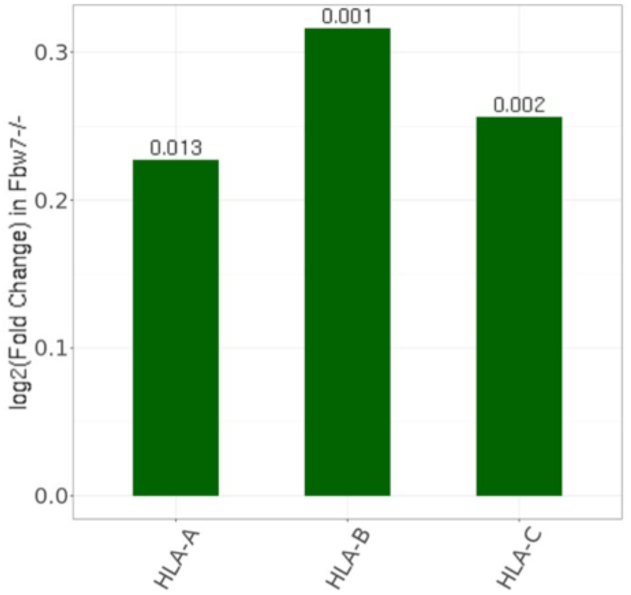
Expression fold change of MHC Class I genes in Hct116 Fbw7^-/-^ with respect to WT cells. FDR values are indicated at top of each bar. n = 3

**Figure 6 – figure supplement 1.**
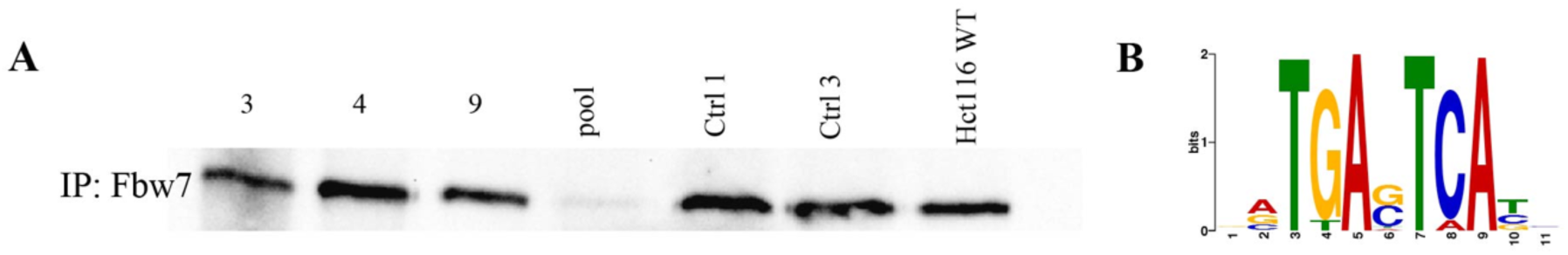
Validation of U5-NSC Fbw7-/- generation and CUT&RUN Jun signal. (A) Western blot showing Fbw7; samples 1-4: U5 NSCs with sgRNA targeting Fbw7 exon 3, 4, 9 and all three exons in one pool; samples 5-6: U5 NSCs with control sgRNA 1x and 3x; and sample 7: Hct116 WT. (B) Sequence logo of AP-1 motif enriched in Jun peaks in U5 NSCs (E value = 1.2e-146).

**Figure 6 – figure supplement 2.**
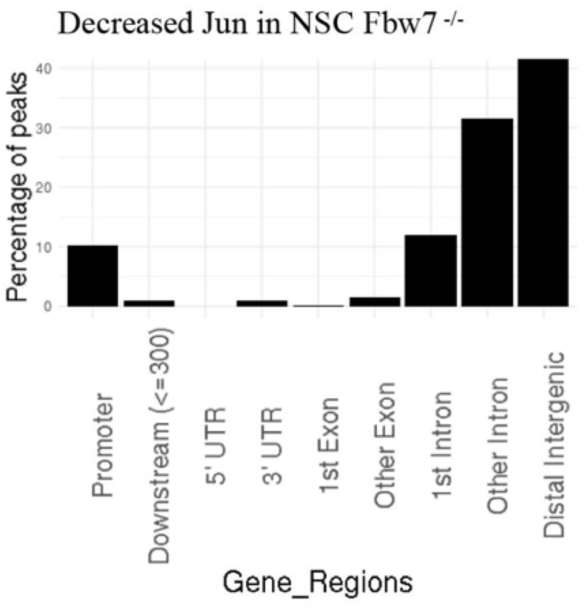
Percentage of peaks with decreased Jun in U5 NSC Fbw7^-/-^ within different gene regions.

**Figure 6 – figure supplement 3.**
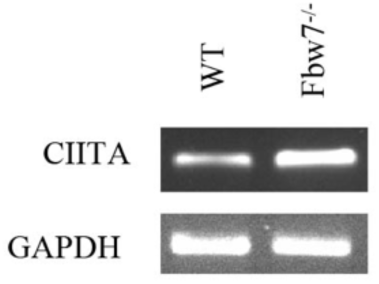
CIITA isoform III amplified using isoform specific primers in U5 NSCs.

